# Evolution of resistance to Irinotecan in cancer cells involves generation of topoisomerase-guided mutations in non-coding genome that reduce the chances of DNA breaks

**DOI:** 10.1101/2021.11.26.470089

**Authors:** Santosh Kumar, Valid Gahramanov, Shivani Patel, Julia Yaglom, Lukasz Kaczmarczyk, Ivan Alexandrov, Gabi Gerlitz, Mali Salmon-Divon, Michael Y. Sherman

## Abstract

Resistance to chemotherapy is a leading cause of treatment failure. Drug-resistance mechanisms involve mutations in specific proteins or changes in their expression levels. It is commonly understood that resistance mutations happen randomly prior to treatment and are selected during the treatment. However, selection of drug-resistant mutants in culture could be achieved by multiple drug exposures of cloned genetically identical cells, and thus cannot result from selection of pre-existent mutations. Accordingly, adaptation must involve generation of mutations de-novo upon drug treatment. Here we explored the origin of resistance mutations to a widely used Top1 inhibitor irinotecan, which triggers DNA breaks, causing cytotoxicity. Resistance mechanism involved gradual accumulation of recurrent mutations in non-coding regions of DNA at Top1-cleavage sites. Surprisingly, cancer cells had higher number of such sites than reference genome, which may define their increased sensitivity to irinotecan. Homologous recombination repair of DNA double strand breaks at these sites following initial drug exposures gradually reverted cleavage-sensitive “cancer” sequences back to cleavage-resistant “normal” sequences. These mutations reduced generation of DNA breaks upon subsequent exposures, thus gradually increasing the drug resistance. Together, large target size for mutations and their Top1-guided generation lead to their gradual and rapid accumulation, synergistically accelerating development of resistance.

**Abstract Figure:** 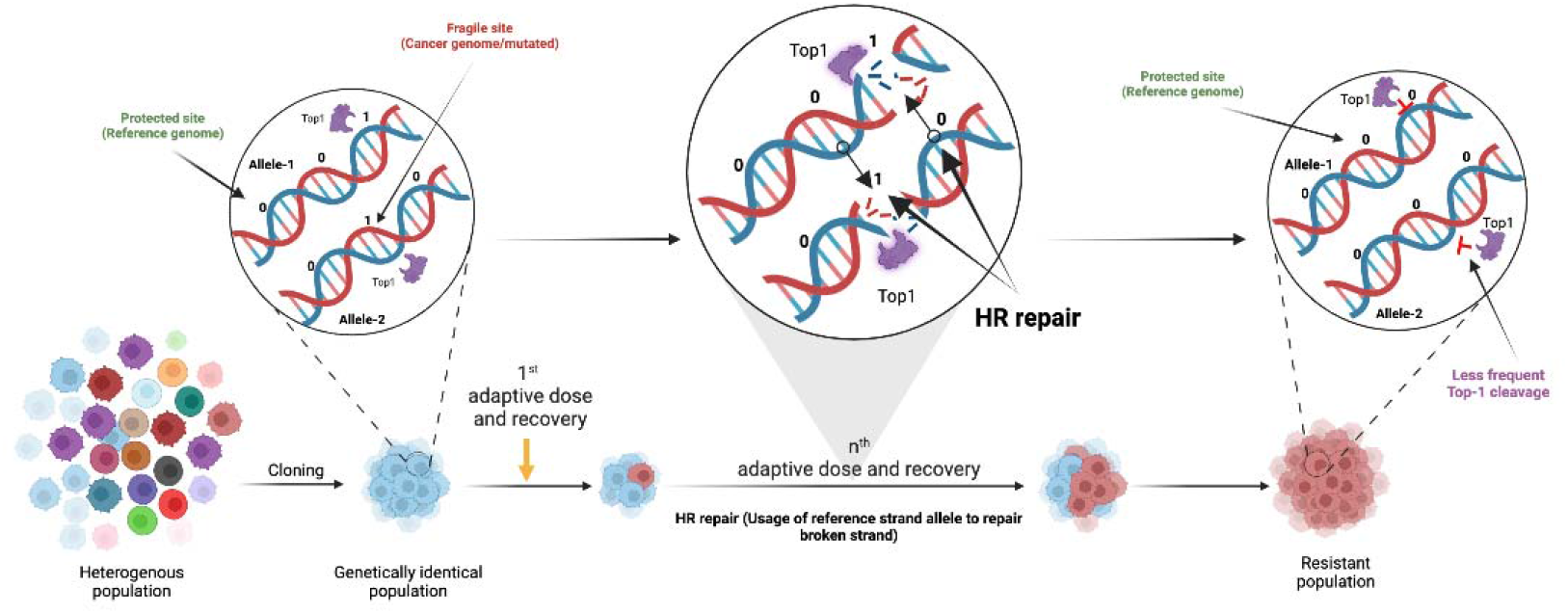

## Background

Resistance to chemotherapy is a leading cause of treatment failure. In line with prior knowledge of development of antibiotic-resistant bacteria (Webster, 1937; Luria & Delbrück, 1943; Cairns *et al*, 1988; Hall, 1988; Kohler *et al*, 1991; Sprouffske *et al*, 2018), the major concept in the cancer field is that resistance mutations happen randomly prior to a treatment and are positively selected during the treatment. This notion was directly supported by experiments with cell barcoding that studied selection of resistant clones followed by the barcode analysis (Research, 2015; Jacobs *et al*, 2020; Jiménez-Sánchez *et al*, 2020; Meyer *et al*, 2020; Patwardhan *et al*, 2021). In these experiments a fraction of clones selected in parallel independent treatments had same barcodes, strongly suggesting that these clones originated from the same parental cells which carried resistant mutations prior to the beginning of drug treatment (Bhang *et al*, 2015; Kress *et al*, 2015; Jahn *et al*, 2018). However, other selected resistant clones that carried different barcodes were not further investigated in these studies and may result from mutations that were generated in response drug treatment. Overall, in selection of drug-resistant mutations in culture, cells are adapted to low drug concentrations, and then via multiple passages with dose escalation, resistant mutants are selected (Demerec, 1937; Cairns, 1975; Hall, 1988; Jackson & Loeb, 1998; Cahill *et al*, 1999; Duesberg *et al*, 2000; Kumar & Subramanian, 2002; Loeb *et al*, 2003; Jensen *et al*, 2015). The process usually takes several months and provides resistance to higher than initial drug concentrations, but not full drug resistance. Importantly, selection of drug-resistant mutants in dose escalation experiments could be achieved from cloned genetically identical cell population, suggesting that adaptation must involve generation of mutations de-novo in the process of drug treatment.

Exploration of forces driving drug-resistance under the dose-escalation, may help to uncover novel mechanisms of development of resistance in clinical setting (Duesberg *et al*, 2000; Pang *et al*, 2013; Sadanandam *et al*, 2013; Jensen *et al*, 2015; Phipps *et al*, 2015; Sinicrope *et al*, 2016; Lee *et al*, 2017; Venook *et al*, 2017; Patnaik & Anupriya, 2019; Raskov *et al*, 2019; Russo *et al*, 2019; Vitiello *et al*, 2021). Indeed, multiple administration of drug doses in clinic appears to mimic the in vitro scheme of drug resistance selection. Understanding the mechanisms of the development of drug resistance can also provide novel insights to guide the design of drug combinations and treatment strategies.

Here, we investigated how these adaptive mutations may emerge in colon cancer cells with the example of resistance to a widely used anti-cancer drug irinotecan. Irinotecan is a pro-drug, which is converted into the active drug SN-38, which binds to Top1 (Champoux, 2001; Vanhoefer *et al*, 2001; Pommier, 2013; Liu *et al*, 2015). The binding, in turn, allows Top1 to make DNA breaks, but prevents re-ligation (Pommier, 2013). Top1-mediated single strand breaks may facilitate double strand DNA breaks that, if unrepaired, cause cancer cell death (Champoux, 2001). Top1 works both during transcription and replication (Pommier, 2013). A number of mechanisms of resistance to Irinotecan have been described, including mutations in Top1, upregulation of multidrug-resistance pumps and associated enzyme system (Vanhoefer *et al*, 2001; Gottesman, 2002; Alfarouk *et al*, 2015; Hammond *et al*, 2016; Hu *et al*, 2016). It is difficult, however, to understand why a dose escalation scheme could be important for development of drug resistance based on these mechanisms. Here, we evaluated the efficiency of the development of irinotecan resistance and uncovered a novel resistance mechanism based on the active generation of a large number of mutations in Top1-dependent DNA breaking sites that reduce the chances of double strand breaks upon consequent irinotecan exposures in the process of dose escalation.

## Methods

### Cell culture and reagents

Cell lines were obtained from the ATCC. HCT116 cells (ATCC Cat# CCL-247, RRID:CVCL_0291) were cultured in McCoy’s 5A Medium supplemented 10% FBS, 4mM L-glutamine (Cat#03-020-1B, BI-Biologicals), 2mM L-alanyl-L-glutamine (Cat#03-022-1B, BI-Biologicals), 1% penstrep (Cat#03-031-1B, BI-Biologicals), and were grown in a humidified incubator at 37°C and 5% CO2 (Meriin *et al*, 2018). SN-38 was purchased from Sigma-Aldrich (MO, USA).

### Single cell line cloning

Single cell cloning was performed by limiting dilution method. Using cloning discs, individual clones were isolated (Merck, Ca#Z374431), grown and stored (10% DMSO in FBS) as stocks in liquid nitrogen for further use.

### Cell cytotoxicity assay

Cells were plated at 30% confluency and treated with the drug at specified concentrations according to experimental plan. Following treatment for a specified period of time, the drug was removed and cells were washed with 1x-PBS, followed by fixation with 1.2% formaldehyde for 10min. After washing, cells were stained with DAPI (1:5000 dilution) for 5min and washed 4 times with 1x-PBST. Imaging was performed using Hermes Wiscon Imaging System (IDEA Bio-Medical Ltd., WiScan Hermes High Content Imaging System, RRID:SCR_021786) and image analysis was performed using inbuilt software package system (Athena Wisoft, Ver1.0.10) “count cell algorithm”. DAPI-positive fixed cells were counted and compared to the untreated control for quantification of drug response as described (Patel *et al*, 2022).

### Virus Preparation for the barcoding library and shRNA screens

Lentiviral libraries for barcoding and shRNA screens were prepared according to the manual. Briefly, cells were passaged and grown at 80-90% confluency. For transfection, reagents plasmids and Lipofectamine 3000 were mixed in opti-MEM (Thermo Scientific, MA, USA) supplemented with 4mM glutamine and co-incubated for overnight. Next day media was changed with OptiMEM supplemented with 5% FBS, 4mM glutamine and kept for 24 hours. Viruses were harvested using 0.45 μm filter and kept in −80C until further use. Upon infection with corresponding libraries, we chose the MOI of about 20%, so that on average each cell receives only one viral particle. After infection, cells carrying lentiviruses were selected with puromycin and further divided into groups for drug treatment in culture(Yaglom *et al*, 2018).

### Cell barcoding

Cloned cell population was barcoded with 50 million (50M) library according to manufacturer protocols (Cellecta CloneTracker 50M Lentiviral Barcode Library, RRID:SCR_021827(Seth *et al*, 2019)). Briefly, cells were infected with the barcoding lentiviruses, selected with puromycin and further divided into groups for drug treatment. After the treatments, cells were allowed to recover, genomic DNA was purified and barcodes were isolated by nested PCR, all samples were multiplexed for sequencing. Detailed procedure of PCR and primers details for 50M libraries is available in supplementary datasheet “Data_S1” (Tables S1 & S2).

### Pooled shRNA Genetic Screen

For the shRNA screen, we used human decipher module 1 library (RRID:Addgene_28289)(Diehl *et al*, 2014; Khorashad *et al*, 2015; D’Alesio *et al*, 2016). Cells were infected with this pooled shRNA library with low MOI. Cells were treated with SN-38 for 24 hours, recovered for 4 days, and collected for DNA isolation and further processing. Barcodes were isolated by nested PCR, sequenced and analyzed, according to the manufacturer’s protocol.

### Genomic DNA extraction and amplification of library barcodes

Isolation of genomic DNA from cultured cells was performed by Wizard genomic DNA isolation kit (Promega, WI, USA). Amplification of the barcodes was carried out by nested PCR. Detailed procedure of PCR and primer details for shRNA screen is available in Supplementary datasheet “Data S1” (Tables S3& S4). Briefly, 1-st PCR (PCR 1) was performed using Titanium Taq DNA Polymerase (# 639209, Takara Bio, CA, USA). Separation of the PCR products from primers and gel purification was done by QIAquick PCR & Gel Cleanup Kit (Qiagen, Germany). 2-nd PCR (PCR 2) was carried out using nested primers either generic or having unique sample barcodes. PCR 2 was performed using Phusion High-Fidelity PCR Master Mix (Thermo Scientific, MA, USA). Samples were multiplexed by adding an additional sample barcode during the second round of PCR. Samples were normalized individually, then pooled together, and purification of the PCR products was completed using AmpureXP magnetic beads (Beckman Coulter, CA, USA) following manufacturer protocols. Next, we sequenced the barcodes using Ion Torrent.

### Analysis of the barcode data

We used a combination of custom-tailored applications to analyze sequencing reads along with the R packages. Data were first checked for quality of reads through FastQC (v0.11.7, RRID:SCR_014583), further using barcode-splitter (v0.18.6, barcode splitter, RRID:SCR_021825) reads were demultiplexed based on sample barcodes (1 error as mismatch or deletion was allowed for sample barcodes while demultiplexing). Obtained FASTQ files were used to count the library barcodes by using python-based applications that were custom-made for this purpose. Quantification of the unique barcodes that were enriched or lost after treatment has been done via a python-based script (software version 3.10.0, RRID:SCR_008394). For data cleaning and visualizations tidyverse-v1.0.0 (RRID:SCR_019186) and ggplot2-v3.3.3 (RRID:SCR_014601)(Wickham, 2016) R packages were utilized.

### Transcriptome analysis

RNA was extracted from cells using the RNeasy Mini kit (Cat#74104, Qiagen). Library preparation strategy (BGISEQ-500, RRID:SCR_017979) was adopted and performed by BGI, China. Briefly, mRNA molecules were purified from total RNA using oligo(dT) attached magnetic beads and fragmented into small pieces using fragmentation reagent after reaction a certain period at proper temperature. First-strand cDNA was generated using random hexamer-primed reverse transcription, followed by a second strand cDNA synthesis. The synthesized cDNA was subjected to end-repair and then was 3’ adenylated. Adapters were ligated to the ends of these 3’ adenylated cDNA fragments. This process amplified the cDNA fragments with adapters from previous step. PCR products were purified with Ampure XP Beads (AGENCOURT), and dissolved in EB solution. Library was validated on the Agilent Technologies 2100 bioanalyzer (2100 Bioanalyzer Instrument, RRID:SCR_018043). The double stranded PCR products were heat denatured and circularized by the splint oligo sequence. The single strand circle DNA (ssCir DNA) were formatted as the final library. The library was amplified with phi29 to make DNA nanoball (DNB) which had more than 300 copies of one molecular. The DNBs were load into the patterned nanoarray and single end 50 (pair end 100) bases reads were generated in the way of sequenced by synthesis. For in house pipeline was developed to analyze the data where reads were first trimmed and clipped for quality control in trim_galore (v0.5.0, RRID:SCR_011847) and checked for each sample using FastQC (v0.11.7, RRID:SCR_014583). Data was aligned by Hisat2 (v2.1.0, RRID:SCR_015530) using hg38, GRch38.97. High-quality reads were then imported into samtools (v1.9 using htslib 1.9, RRID:SCR_002105) for conversion into SAM files and later to BAM file. Gene-count summaries were generated with featureCounts (v1.6.3, RRID:SCR_012919): A numeric matrix of raw read counts was generated, with genes in rows and samples in columns, and used for differential gene expression analysis with the Bioconductor, RRID:SCR_006442 (edgeR v3.32.1, RRID:SCR_012802 (Chen *et al*, 2021), Limma v3.46.0, RRID:SCR_010943) packages to calculate differential expression of genes. For normalization “voom” function was used, followed by eBayes and decideTests functions to compute differential expression of genes.

### Human Whole Genome (HWG) sequencing and analysis

Isolated DNA was sent for 30X whole genome sequencing as a service by commercial provider Dante labs, Italy. After obtaining reads, data were first checked for quality of reads through FastQC (v0.11.7, RRID:SCR_014583), further using barcode-splitter (v0.18.6) reads were demultiplexed based on sample barcodes (1 error as mismatch or deletion was allowed for sample barcodes while demultiplexing). Reads were aligned to reference genome hg38 with BWA-MEM (RRID:SCR_010910)(Li & Durbin, 2010) default settings. Aligned SAM files were converted to BAM using samtools (ver1.9, RRID:SCR_002105). Obtained BAM files were used to call variants using Genome Analysis Tool Kit (GATK v4.1.8.0, RRID:SCR_001876)(Geraldine A. Van der Auwera, Brian D. O’Connor, 2020) from Broad’s Institute, we called for combined genomic variants as Variant Call Factor (VCF) file as output. In above analysis we followed strictly the best practices guidelines of GATK(DePristo *et al*, 2011). All further analysis was done using either suitable R packages (tidyverse-v1.0.0 (RRID:SCR_019186) and ggplot2-v3.3.3(RRID:SCR_014601), karyoploteR v1.18.0(karyoploteR, RRID:SCR_021824)(Gel & Serra, 2017) for data, we also used SnpEff (v3.11, RRID:SCR_005191)(Cingolani *et al*, 2012) program for annotation of genes and bedtools v2.27.1(RRID:SCR_006646)(Quinlan & Hall, 2010) to match obtained variants to available bed files from UCSC(UCSC Cancer Genomics Browser, RRID:SCR_011796)(Kent *et al*, 2002), GEO (Gene Expression Omnibus (GEO), RRID:SCR_005012) datasets and other relevant databases. To extract data from metadata file and for comparative analysis we used Microsoft excel (RRID:SCR_016137) as tool. Raw data files and associated common mutation analysis in form of VCF file is available at SRA (NCBI Sequence Read Archive (SRA), RRID:SCR_004891) under accession number (PRJNA738674).

### Assessment of cluster mutations in resistant mutants

For estimating the number of mutations present in close proximity (clusters), we took 100bp small windows and calculated whether the mutations were present in close proximity in these small intervals. We used custom written python codes to first, separate the whole genome into small bins of 100bp respectively. Further we defined common mutation’s genomic locations to it and calculated the conditions that is how many of these mutations fall within 100bp window. The output was generated as text file where genomic locations and number of mutations in 100bp window were calculated.

### Classification of repeats and computing mutations in low complexity regions of genome

For classifying the nature of repeats where these mutations have been accumulated, we used RepeatMasker v4.1.0 (RepeatMasker, RRID:SCR_012954, database Dfam_3.1 (RRID:SCR_021168) and rmblastn version 2.9.0)(Smit, AFA, Hubley, R & Green, P) that screens DNA sequences for known repeats and low complexity DNA sequences. Reference sequence hg38 was used for this purpose. Common (triple) mutations (total-46099) were aligned to reference database according to their occurrences based on genomic locations.

Program defines the nature of the sequence where these mutations have been accumulated. The output of the program is a detailed annotation of repeats that are present in the query sequence as well as a modified version of the query sequence in which all the annotated repeats are classified. Sequence comparisons were performed on Unix environment using cross-match, an efficient implementation of the Smith-Waterman-Gotoh algorithm developed by Phil Green, or by WU-Blast developed by Warren Gish (Smit, AFA, Hubley, R & Green, P). Output file is matrix of repeated elements found in query sequence compared to reference libraries reported as percentage in the tabulated form (Flynn *et al*, 2019).

### Assessment of satellites and simple repeats

In-depth analysis was done to identify the nature of satellites and its classification using RepeatMasker v4.1.0 (RepeatMasker, RRID:SCR_012954) (Smit, AFA, Hubley, R & Green, P) advanced options wherein we investigated the type of satellites and simple repeats. Firstly, fasta sequences for the genomic locations for common mutations were fetched using bedtools “getfasta” algorithm (BEDTools, RRID:SCR_006646)(Quinlan & Hall, 2010). It was further analyzed by RepeatMasker v4.1.0(Smit, AFA, Hubley, R & Green, P) to identify the repeated elements. Next, “fasta.out” was used to further investigate and quantify the classification of the satellites type. While “fasta.tbl” was used to compute overall percent representation of the repeated elements. Additionally, we used Tandem repeat finder (RRID:SCR_005659) (Benson, 1999) and UCSC microsatellite track to investigate and compute the microsatellites and simple repeats.

### Double strand break quantification

Estimation of double strand break occurring in the system of wild type cell population and resistant lines once adapted to drug was done using (a) γH2AX assay(Ivashkevich *et al*, 2011) and (b) Comet assay(Neri *et al*, 2015).

### γ-H2AX assay

SSC7 parental cells and cells that underwent five cycles of drug exposure (T5_resistant), were exposed to 4nM SN-38 for 24 hours. Alternatively, SSC1 and MSC1, MSC2 & MSC3 cells were exposed to 40nM SN-38 for 30min. Cells were washed with 1xPBS and fixed with 0.2% formaldehyde. Permeabilization was done by 0.2% Triton X-100 in PBS for 10 minutes at room temperature. Then cells were blocked with Bovine serum albumin (BSA, 5% w/v in Phosphate buffered saline with 0.05% tween-20: PBST) for 1 hour. Washed 3 times with PBST. Incubation with primary antibody (Phospho-Histone H2A.X (Ser139) Antibody, Cell Signaling Technology Cat# 2577, RRID: AB_2118010) was done overnight at 4^0^C. Cells were washed with 1xPBST five times followed by incubation with secondary antibody (Goat Anti-Rabbit IgG H&L, Alexa Fluor® 488, Abcam Cat# ab150077, RRID:AB_2630356) for 1 hour. Antibody was removed and washed five times to remove non-specific antibody binding. Imaging was performed using Hermes Wiscon Imaging System (IDEA Bio-Medical Ltd., WiScan Hermes High Content Imaging System, RRID:SCR_021786) and image analysis was performed using inbuilt software package system (Athena Wisoft, Ver1.0.10). The software takes the maxima and minima for the foci intensity. For all the analysis we kept constant maxima=550 to select the foci and selected automatic background correction based on its untreated sample. Statistics calculation was performed with GraphPad Prism version 9.0.0 (GraphPad Prism, RRID:SCR_002798), California USA. The significance of differences was determined using unpaired Welch’s correction, two-tailed t-test (“ns” <0.1234, *P < 0.0332, **P < 0.0021, ***P < 0.0002, ****P <0.0001).

### Single cell gel electrophoresis (SCGE-Alkaline)

To estimate the change in break of DNA strands before and after treatment in normal cells and resistant cell population, a single cell gel electrophoresis (SCGE) or COMET assay was performed(Singh *et al*, 1988). Cells were embedded in 1% agarose on a microscope slide are lysed with detergent and high salt.

Glass slides were precoated with 1% agarose. Cells were treated with SN-38 or left untreated to serve as control. Preparation of sample was done by scrapping cells gently with 0.05% trypsin. Washed 3 times with ice cold 1X-PBS and roughly 0.1*10^6^ cells/ml was taken. 50uL of cell suspension and 50uL of 1% low molten agarose kept at 37^0^C was mixed together. 75uL was used to make bubbles on the slide and was left to solidify at 4^0^C until it formed clear ring. Lysis was performed by placing the sample slides in lysis buffer [(Nacl (2.5M), EDTA-pH 10 (100mM), Tris-Base pH 10 (10mM), Triton X100 (1% freshly added before use), buffer pH was maintained at 10 before adding Triton-X100] overnight at 4^0^C. Unwinding was performed by rinsing the slides with fresh water to remove salts and detergents. Slides were immerged in unwinding buffer [(NaOH(300mM), EDTA (1mM), buffer pH 13)] for 1 hour at 4^0^C. After unwinding, samples were run in electrophoresis alkali buffer [(NaOH (12g/L), EDTA (500mM pH 8)], slides were kept in electrophoresis tank and buffer was poured to cover the slides. Running was performed at 22V(constant), 400mA(constant) for 40 minutes. Neutralization was performed by dipping the slides in tris buffer (0.4M, pH 7.5) for 5 minutes. Slides were then immerged in 70% ethanol for 15 minutes, air dried for 30 minutes, Staining was performed by incubating in DAPI (1ug/ul, 1:5000 dilution). Washed with ice cold water and air dried in dark. Imaging was performed using Olympus IX81 Inverted Fluorescence Automated Live Cell Microscope (18MP CMOS USB camera) with associated, in-built software package (Olympus IX 81 Inverted Fluorescence Automated Live Cell Microscope, RRID:SCR_020341). Quantification was performed using OpenComet v1.3.1(OpenComet, RRID:SCR_021826)(Gyori *et al*, 2014). Statistics calculation was performed with GraphPad Prism version 9.0.0 (GraphPad Prism, RRID:SCR_002798), California USA. The significance of differences was determined using unpaired Welch’s correction, two-tailed t-test (*P < 0.0332, **P < 0.0021, ***P < 0.0002, ****P <0.0001).

### Cell cycle reporter assay

Reporter plasmid (pBOB-EF1-FastFUCCI-Puro, Addgene-plasmid # 86849; http://n2t.net/addgene:86849; RRID:Addgene_86849) was made by Kevin Brindle & Duncan Jodrell (Koh *et al*, 2017). Lentiviruses were produced in the lab as described in Materials and Method section above. Cells were infected with the viruses and selected with puromycin. Treatment was carried out with SN-38(4nM) for 24 hours. After the drug removal, recovery was recorded by live imaging on alternate days. “Control” is referred as cells before treatment, and images were taken at consecutive days till 9^th^ day of recovery. Analysis was performed using FIJI-ImageJ program using imaged channels.

### DNA-Top1 adduct capturing RADAR assay

HCT116 (SCC1 parental clone) and mutant SCC1 (MSC1 and MSC2) cell lines were seeded in 6 well plates in 2 ml of McCoy’s 5A Medium supplemented 10% FBS, 4mM L-glutamine (Cat#03-020-1B, BI-Biologicals), 2mM L-alanyl-L-glutamine (Cat#03-022-1B, BI-Biologicals), 1% penstrep (Cat#03-031-1B, BI-Biologicals) at 25% of confluence and were cultured overnight in a humidified incubator at 37°C and 5% CO2. The next day the media was aspirated, the cells were washed with 1xPBS and treated with 40nM irinotecan for 6 hours. After treatment, medium was removed and cells were lysed on the plate by addition of 3 ml of lysis reagent RLT Plus (Lot 169010027, Qiagen, Germany). 1 ml of lysate was transferred to a 2 ml Eppendorf tube, then 0.5 ml of 100% ethanol (1/2 volume) was added.

The mix was incubated at −20°C for 5 min, and centrifuged for 15 min at 16000 rcf. Another volume of lysate was frozen and stored in −80°C. After centrifugation the supernatant was aspirated and the pellet containing nucleic acids in complex with proteins was washed twice in 1 ml of 75% ethanol by vortexing followed 10 min centrifugation. Finally, the pellet was diluted 20 μl of 8 mM NaOH and 20 μl of Tris-Buffered Saline buffer (150 mM NaCl, 50 mM TrisHCl, pH 7.6: TBS). The quantity of DNA was measured with Thermo Scientific Nanodrop spectrophotometer, then the samples were normalized with TBS buffer. The small volumes 5 μl of solubilized DPCC isolates were transferred to a nitrocellulose membrane as dots. After the samples were dried the blocking procedure with Bovine serum albumin (BSA, 3% w/v in Phosphate buffered saline with 0.05% tween-20: PBST) followed for 1 hour in room temperature. After blocking the membrane was washed with PBST 3 times for 5 min on shaker, then incubated with 10 ml of the primary antibody detecting TOP1 (Invitrogen) (1:1000 in 1.5% BSA in PBST) for overnight in 4°C on shaker. After incubation the membrane was washed with PBST 3 times for 5 min on shaker, 10 ml of the secondary antibody Peroxidase-AffiniPure Goat Anti-Rabbit IgG (H+L) (Cat#111-035-003, Jackson Immuno Research Inc, USA) (1:3000 in 1.5% BSA in PBST) was added and incubated for 1 hour in room temperature on shaker. Before the detection the membrane was washed with PBST 5 times for 5 min on shaker, after 1 ml of Immobilon Forte Western HRP Substrate (Cat# WBLUF0500, Millipore, Germany) was added and incubated with shaking for 1 minutes. The detection was performed with BIORAD Protein detection system. Quantification was performed on Fiji image-J. Statistics calculation was performed with GraphPad Prism version 9.0.0 (GraphPad Prism, RRID:SCR_002798), California USA. The significance of differences was determined using unpaired Welch’s correction, two-tailed t-test (“ns” <0.1234, *P < 0.0332, **P < 0.0021, ***P < 0.0002, ****P <0.0001).

### Immunoprecipitation and Immunoblotting

Cells were lysed with lysis buffer (50mM Tris-HCl (pH 7.4), 150mM NaCl, 1% Triton X-100, 5mM EDTA, 1mM Na3VO4, 50mM β-glycerophosphate, 50mM NaF) supplemented with Protease Inhibitor Cocktail (Cat. #P8340, Sigma) and Phenylmethylsulfonyl fluoride (PMSF). Samples were adjusted to have equal concentration of total protein and subjected to PAGE electrophoresis followed by immunoblotting with primary antibody (Phospho-Histone H2A.X (Ser139) Antibody, Cell Signaling Technology Cat# 2577, RRID:AB_2118010)(Meriin *et al*, 2018).

### Hypergeometric distribution

To predict the significance of overlap between triple mutations and Top1-cleavage sites we used R language tool (see code availability section for applicable codes). The problem of DNA sites overlap is described by a hypergeometric distribution where one list defines the number of cleavage sites and the other list defines the number of mutations. Assume the total genome size is *n*, the number of points in the first list is *a,* and the number of points in the second list is *b*. If the intersection between the two lists is *t*, the probability density of seeing *t* can be calculated as: *Probability of occurrence=dhyper (t, a, n – a, b).* Based on the existing data, *b* < *a*. So, the largest possible value for *t* is *b*. Therefore, the p-value of seeing intersection *t* is: *sum (dhyper (t:b, a, n - a, b))*. Taking, top1 cleavage sites (a) cover 2.5*10^7^ base pairs in the genome (2*10^5^ sites X 50bp per peak), mutations (b)=46099, total genome (n)=3*10^9^ bp, t=9000 overlap out of 46099. Sum of overlap was computed to be absolute zero with null probability.

## Results

### Experimental design with multiple irinotecan (SN-38) treatments

Since irinotecan is widely used against colon cancer, we investigated its effects on colon cancer cell line HCT116. Since unlike in the organism, in cell culture, irinotecan is activated very ineffectively, in all experiments below we used already activated derivative of irinotecan i.e. SN-38. In order to achieve genetic uniformity of the population, we cloned the cells and isolated several independent clones. Genetic uniformity within each clone indicates that any drug-resistance mutation selected in our experiments occurs either in the process of the colony growth from a single cell prior to the drug treatment or is actively generated in the process of drug treatment. Sensitivity to SN-38 differed dramatically between the clones, with minimal toxic concentration between 1nM and 80nM. (Figure S1). We chose two clones, SSC1 and SSC7, with high sensitivity (IC_50_ 1nM and 2nM, respectively) for further experiments.

To understand development of drug resistance, SSC7 cells were exposed to 4nM SN-38, which led to the death of a significant fraction of the population, and the cell cycle arrest of the rest of the population. Cells remained in the arrested state without divisions for 14 days, and then resumed growth. The arrest was associated with a senescence-like phenotype (highly enlarged and vacuolized cells. Second treatment with 4nM SN-38 led to a shorter period of the cell cycle arrest. Such a treatment cycle was conducted five times in total. At the fifth cycle, practically no cell death or growth inhibition was seen, indicating that cells became fully adapted to this concentration of SN-38 (Figure 1A, B).

**Figure 1.**
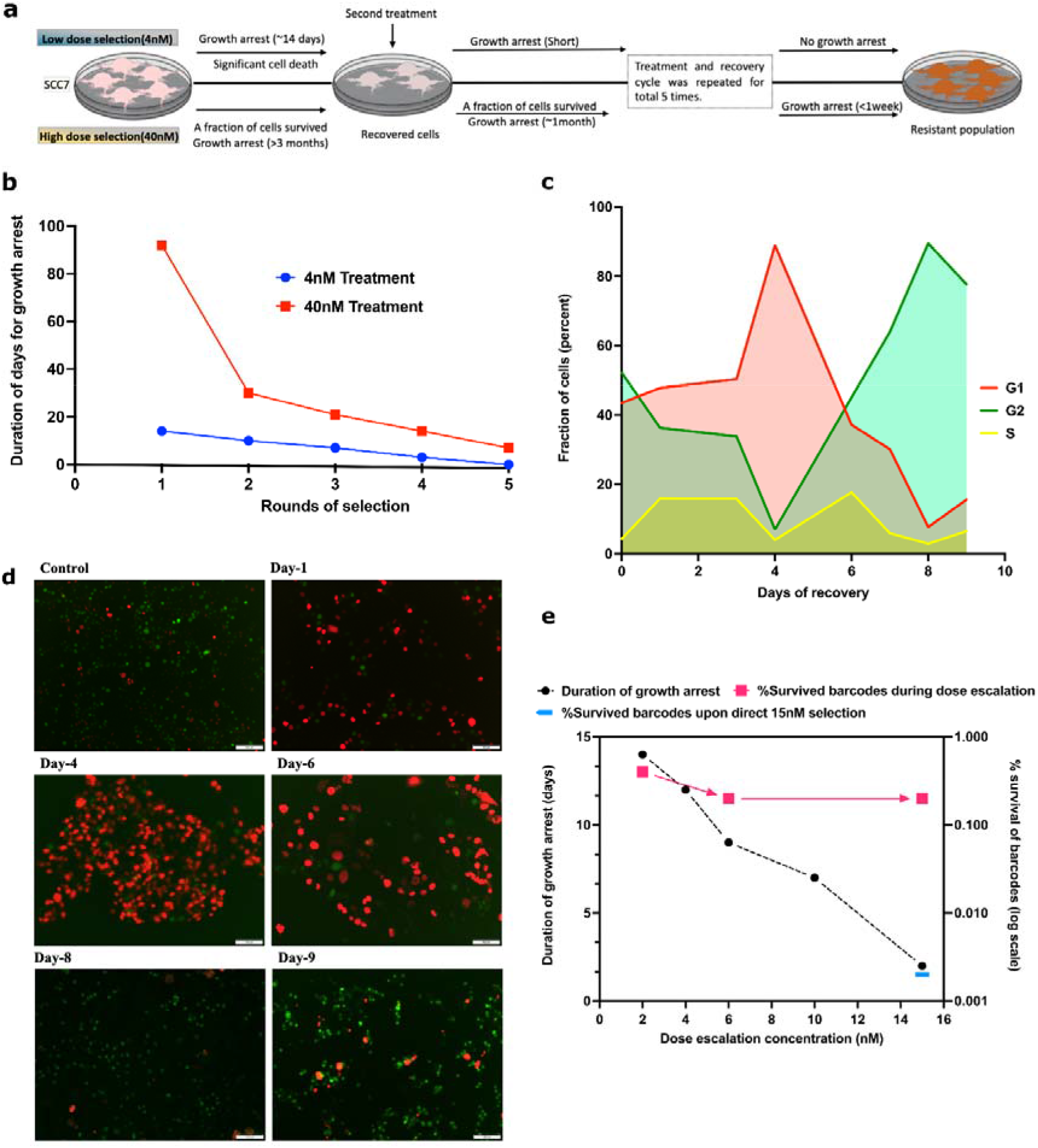
Adaptation to SN-38 associates with reduction in duration of growth arrest: **(a)** Graphical sketch of the experiment plan, showing low dose selection (upper panel), high dose selection (lower panel). **(b)** clone of HCT116 cells was exposed to the same dose of SN-38 (either 4nM or 40nM for 24 hours) multiple times, and periods of the cell cycle arrest following exposures were measured. **(c, d)** Simultaneous recovery of SN-38 treated cells from the cell cycle arrest. Cells were infected with cell cycle reporter plasmid (see Materials and Methods). Cells were treated with 4nM SN-38, and following the cell cycle arrest (Day 1 and 4) allowed to recover (Day 8). Imaging was performed at the indicated days following SN-38 exposure using TexRed (red) and FITC (green) channels. Green fluorescence represents G2, Red fluorescence represents G1 and overlapping of red-green shows S phase. Quantification of data for fraction of G1, G2 and S phase cells during recovery from 4nM SN-38 treatment was quantified by imageJ, experiment was done in triplicate wells for each treatment. **(e)** Cells were exposed to rounds of treatments with 2, 4, 6, 10 and 15nM SN-38 and the periods of recovery were measured. On the same graph survival of barcodes after these treatments is shown. Cells were barcoded using the 50 million lentiviral barcoding libraries (Cellecta), and cells recovered after the treatments were collected, barcodes isolated, sequenced by IonTorrent and analyzed. Quantification of barcode survival is represented on right Y axis. 0.4% of total barcodes survived 2nM dose compared to untreated control (100%). Following further 6nM and 15nM 0.2% of initial barcodes survived, i.e., approximately 50% of barcodes that survived the 2nM dose. Also shown the fraction of barcodes that survived 15nM dose without preadaptation to lower doses (0.002%).

Further, we tested if a similar pattern occurs when cells develop adaptation to high doses (40nM) of SN-38. The fraction of cells that survived 40nM stayed in a senescence-like state for more than three months, after which cells resumed propagation and filled the plate. Upon subsequent exposure of the recovered population to 40nM SN-38 the period of the growth arrest was only about one month. The process was repeated three more times, and each time the fraction of dying cells was lower and the time period of growth arrest was shorter compared to the previous round of selection, so that following the fourth exposure, cells spent approximately one week in the arrested state (Figure 1A, B). Altogether, these findings indicate similarities of adaptation to low and high doses of SN-38.

### Most of the survived cells recover from the senescence like growth arrest

A large fraction of cells in the population could adapt to the treatment and resume growth after the cell cycle arrest. Alternatively, a small fraction of cells that could be originally resistant to the drug continued to propagate, which became detectable only when they began outgrowing the rest of the arrested population. To distinguish between these possibilities, SSC7 cells were infected with the cell cycle reporter virus (Koh *et al*, 2017) and exposed to 4nM SN-38. Survived treated cells stopped dividing, acquired senescence-like phenotype (enlarged, vacuolized cells), and according to the reporter, underwent G1 growth arrest (Figure 1C, D). Cells remained in G1 for four days, after which a majority of cells exited G1 and entered the cell cycle (Figure 1C, D). Entering the cycle following the senescence-like arrest was surprisingly slow, and only by day 8 almost 100% of cells reached G2. There were no localized shifts of cells to G2, indicating the lack of clonal expansion. This observation indicates that cells underwent true adaptation to SN-38, rather than reflects the expansion of a small fraction of initially resistant cells.

### A large fraction of cells gets adapted to SN-38

To quantitatively assess the process of adaptation, 10 million SSC1 cells were individually barcoded using Cellecta 50M barcodes lentiviral library(Seth *et al*, 2019). When cells filled the plate, half of them were collected for the barcode analysis, and half were used for further dose-escalation experiments with sequential passaging over 2, 4, 6, 8, and 15nM of SN-38. With each passage, we observed a reduction of the population of dying cells, and a reduction of the period of growth arrest (Figure 1E). At different stages of the experiment, we collected cells for the analysis of barcodes, as described in Materials and Methods.

Comparison of barcodes that were detected in the control population and population treated with 2nM showed that about 4**\***10^−3^ of the original clones survived the selection (Figure 1E). Analysis of barcodes in the population of cells after the 15nM SN-38 selection demonstrated that 2**\***10^−3^ of original clones survived the entire series of selections, which is only two times lower than the number of clones survived the first round of selection at 2nM. These findings indicate that (a) the probability of the survival of clones is high compared to the usual probability of spontaneous or even mutagen-induced mutations (according to classical studies usually not higher than 10^−4^(Demerec, 1937; Luria & Delbrück, 1943; Cairns, 1975; Kohler *et al*, 1991; Drake *et al*, 1998; Cahill *et al*, 1999; Drake, 1999; Duesberg *et al*, 2000; Kumar & Subramanian, 2002; Loeb *et al*, 2003)), and (b) almost 50% of clones that survive 2nM selection survived the entire series of selections, suggesting that if cells are able to survive initial treatment they can also survive the dose escalation treatment.

To test if the dose escalation process is critical for the development of the resistant forms, barcoded SSC1 cells were exposed directly to 15nM SN-38. Analysis of barcodes indicated that only 2**\***10^−5^ of clones survived (Figure 1E), which is 100X lower than the survival rate of 15nM SN-38 in the dose escalation experiment. These data indicate that the dose escalation protocol is important for effective development of resistant variants.

### Changes in transcriptome may not be involved in the resistance development

To explore the mechanisms of the adaptation, the survived population of multiple rounds of treatment with 40nM SN-38, was cloned again. Barcodes from several clones were isolated and sequenced. Three clones with different barcodes (MSC1, MSC2 and MSC3) were chosen for further analysis. Since the barcoding of cells was done prior to the entire series of SN-38 treatments, the fact that these clones carry different barcodes indicated that they did not split from the same clone somewhere in the middle of the SN-38 treatment. In other words, they represent the progeny of cells that underwent the entire series of the treatments independently of each other. We would like to reiterate that the original barcoded population was genetically homogenous because of the initial cloning. Notably, the selected clones had growth rate similar to the parental SSC1 clone (Figure S2).

To uncover potential mechanisms related to changes in gene expression, we compared transcriptomes of parental SSC1 clone and individual mutant isolates MSC1, MSC2 and MSC3 both untreated or exposed to 10nM SN-38, Experiment was performed in biological duplicates (n=2) for each samples. In naïve condition, a number of differentially expressed genes were observed in the survived clones (Table S5 to Table S10). Importantly, we did not observe changes in either MDR1 or other ABC transporters involved in drug efflux, or Top1, or DNA repair genes, suggesting that the mechanism of resistance in these clones may not be related to expected changes in the transcriptome. Further, datasets were analyzed for the enrichment of genes belonging to known pathways using GSEA. Pathways such as epithelial to mesenchymal transition (MSC2 compared to SCC1) and inflammatory response pathway (MSC2 and MSC3 compared to SCC1) were common. Rest did not qualify FDR threshold of 0.05 in all above comparisons (Table S11), clearly suggesting transcriptome is unlikely to be involved in resistance in the dose-escalation setting.

### DSBs significantly contribute to SN-38-induced cell death

To further explore potential mechanisms of the resistance, we performed a pooled shRNA screen to identify genes important for survival of SN-38 treatment. SSC1 cells were infected with focused lentiviral shRNA library Decipher Module-1(Castel & Martienssen, 2013; Diehl *et al*, 2014; Khorashad *et al*, 2015; D’Alesio *et al*, 2016; Schuster *et al*, 2019) that targets signaling pathways. This library covers about 20% of human genes. Cells were treated with 10nM SN-38 for 24 hours and cells that survived the treatment after 5 days were collected and the barcodes isolated, sequenced and analyzed. As control, we used the same population of infected cells but without SN-38 treatment. Among genes, depletion of which showed sensitizing effects (Table S12), was a group of genes that plays a role in DNA double strand breaks repair, predominantly representing the homologous recombination (HR) and nonhomologous end joining (NHEJ) DNA repair pathways, including POLE, POLE3, POLE4, KAT5, RAD51C, RAD54L, RAD1, RAD9A, H2AFX, LIG4 and PARP2 (Table S13). We also observed a number of genes involved in translesion DNA synthesis (Table S13). The overall list of hits in the screen is shown in table (Table S12). The list of pathways involved in the sensitivity according to GSEA analysis of 1320 hits is shown in Table S14, (Figure 2) where Hallmark DNA repair pathway, Oxidative phosphorylation, Myc, Reactive oxygen species are in top-10 highly enriched pathways. These data reinforce the understanding that (a) generation of double strand DNA breaks is critical for cell death caused by SN-38 and (b) HR and to some extent NHEJ repair pathways play an important role in the SN-38 survival in naïve cells (though these pathways are not significantly upregulated in the resistant clones (Table S11)).

**Figure 2.**
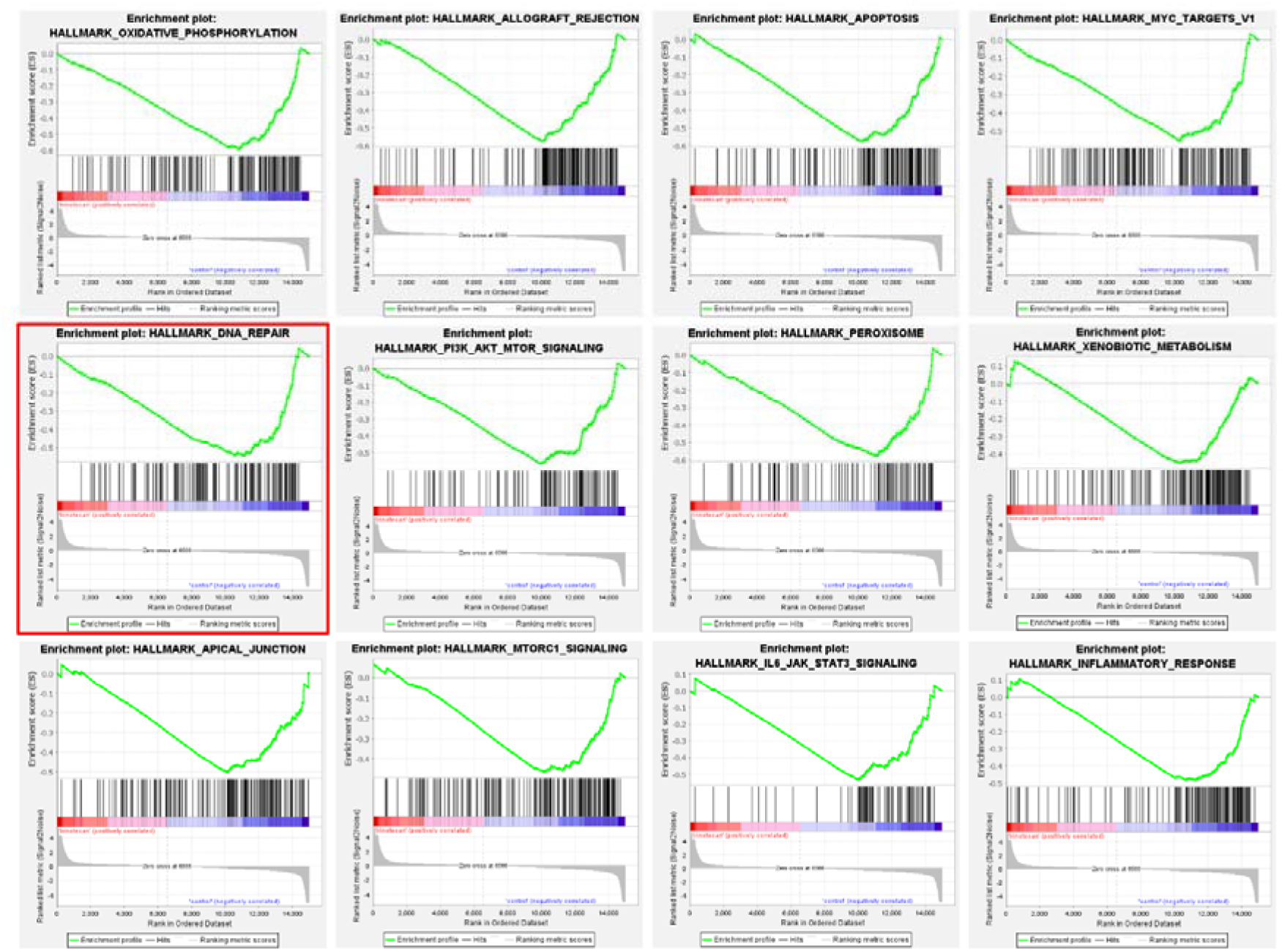
Top-10 enriched pathways uncovered from shRNA screen. A representative plot obtained from GSEA analysis exhibiting important sensitizer pathways (neg-regulated) in response to SN-38, **Top horizontal panel:** Showing Oxidative phosphorylation, Allograft rejection, Apoptosis, Myc. **Middle horizontal panel:** DNA repair (important for survival considering drug favors DNA breaks), MTOR, Peroxisome, and Xenobiotic metabolism (includes MDR’s and drug metabolism related genes), **Lower horizontal panel:** Apical junction, MTORC1, IL6_JAK_STAT3, Inflammatory response. These pathways are top-12 pathways with highly significant False discovery rate (FDR).

### Development of SN-38 resistance associates with emergence of multiple non-random mutations

To study mutations that emerged in the survived clones, we performed whole genome sequencing. Genomes of the survived clones MSC1, MSC2 and MSC3 were compared with the genome of original population SSC1, and each of them with the Reference Genome (GRCh38/hg38) in the UCSC (Kent *et al*, 2002) database. We observed that the parental SSC1 genome had hundreds of thousands of SNPs and InDels compared to the Reference Genome.

These mutations may reflect the fact that HCT116 cells were isolated from a different individual than a group of individuals whose sequence compose the Reference Genome. Alternatively, these mutations could arise in the process of cancer development and/or further culturing of HCT116 cells in the laboratory conditions. Overall analysis of mutation in SSC1 indicated that 93% of them do not correspond to known SNPs in the human population, suggesting that the vast majority of the mutations simply reflect either the cancer nature of these cells or genetic instability upon culturing. Accordingly, they will be further called “cancer alleles”. Notably, HCT116 cells carry a mutation in MLH1 gene that leads to the microsatellite instability(Vilkin *et al*, 2009; Chen *et al*, 2017; Gazzoli *et al*), which may significantly contribute to generation of these cancer alleles, see below. When genomes of the SN-38-resistant isolates MSC1, MSC2 and MSC3 were compared with the genome of SSC1, we identified hundreds of thousands of InDels and SNPs that arose in the process of adaptation to SN-38.

Strikingly a very large fraction of these mutations were common between the independently isolated clones (Figure 3A, B). If compared by pairs, i.e. MSC1/MSC2, MSC1/MSC3 and MSC2/MSC3, in each pair between 17% and 45% of mutations were common, and between 7% and 15% were common between all three independent isolates (46,099 mutations, of which 28.4% were InDels and 71.6% were point mutations (Table S15)). Even considering that SN-38 may have a high mutagenic activity and triggers protective mutations with the rate as high as 10^−4^ per nucleotide, the probability of overlap of a mutation in three independent clones will be 10^−12^ (i.e. way less than one triple mutation per 3*10^9^ bp genome), which is many orders of magnitude lower than seen in the experiment. Thus, the overlapping (common between three clones) mutations clearly point to a non-random mutation mechanism. Generation of these mutations seem to be guided by a mechanism, understanding of which may clarify the adaptation pathway.

**Figure 3.**
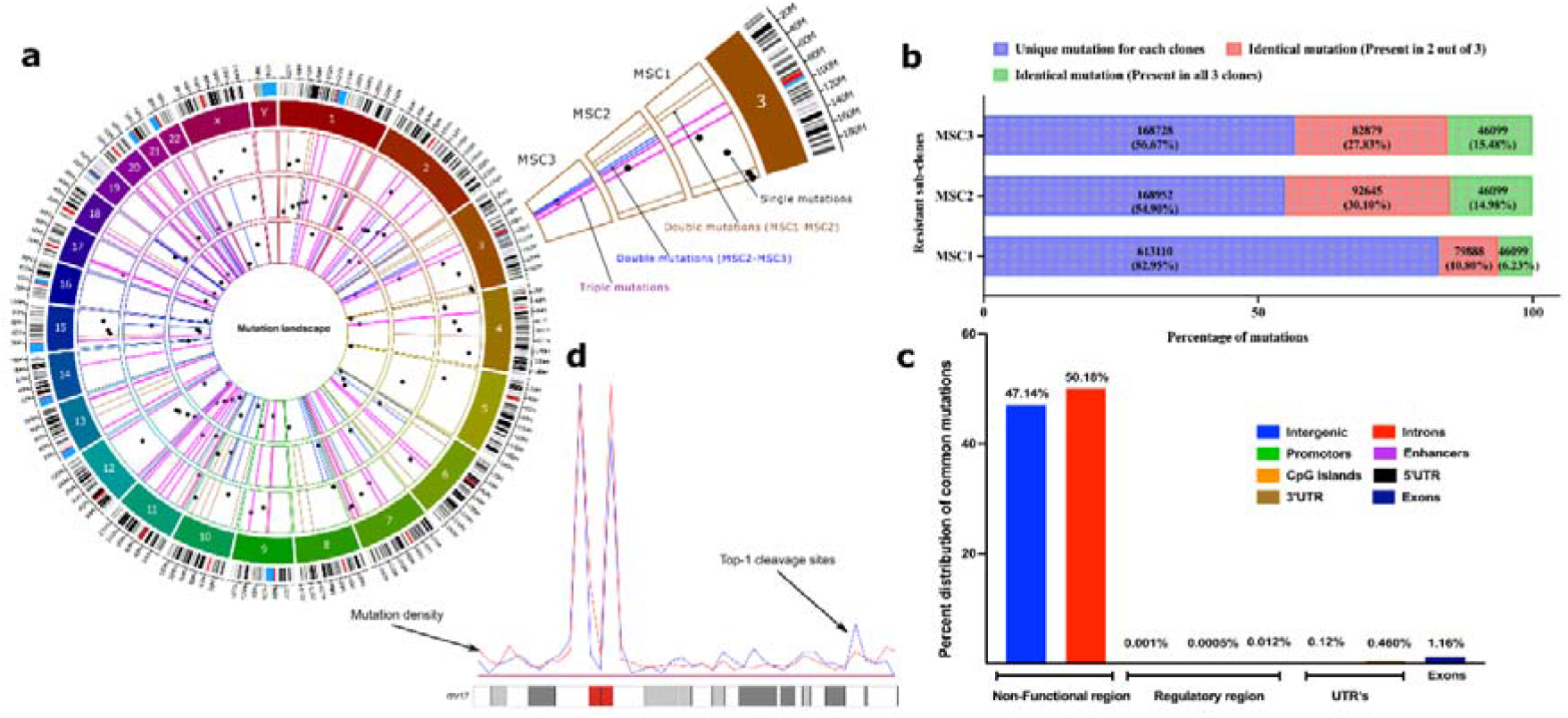
SN-38 resistant subclones harbors identical mutations in genome at non-coding regions: **(a)** Circos plot to represent the category of mutations based on their occurrence in three independent clones. Three circles inside the chromatogram notations represent three individual selected resistant clones (MSC1, MSC2 and MSC3 from outside to inside). Dots (black) inside the respective circles represent mutations that are unique in the clones and not overlapping to others. Bars (brown) represent the overlapping mutations in clone MSC1 and MSC2. Bars (blue) represent the overlapping mutations in clone MSC2 and MSC3. Bars (pink) represent mutations that are overlapping in all three clones. To plot, only 50 mutations of each category was chosen randomly and are plotted for visualization purpose) to their genomic locations. Plot was prepared using commercial license from OMGenomics (Circa, RRID:SCR_021828) application. **(b)** Graph represent the total quantification of overlapping mutations in the SN-38-resistant clones that emerge in the adaptation process. Stack graph showing the overlapping mutations among resistant sub-clones. Mutations in the resistant clones that differ them from SSC1 were compared to each other. These mutations represent three categories; (blue) mutations that are unique to each clone and not found in other clones, (red) mutations that are identical in two clones, (green) mutations that are identical in all three clones, **(c)** Categorical distribution of mutations to the non-functional, regulatory and UTR’s region of genome computed from annotation usingSnpEff, **(d)** An example representation of chromosome-17 showing overlapped mutations density (blue line) and Top-1 cleavage sites (red line).

Analysis of the ENCODE (promotors and enhancer datasets (Libbrecht *et al*, 2019)), GEO (GSE57628, mapping of Top1 sites in human HCT116 cells)(Baranello *et al*, 2016) and UCSC(Kent *et al*, 2002) database datasets uncovered that the vast majority of the mutations are present in heterochromatin (high content of histone H3K9me3) in the noncoding regions (Figure 3C). There was a small fraction of mutations present in promotors/enhancers (<1%) and the coding regions (2%), most of them in the exons (Figure. 3B, C). Importantly, the genes that have mutations in their coding and regulatory regions did not belong to known pathways associated with either cell survival or DNA repair, and therefore are unlikely to be involved in the adaptation process (Table S16, also see mutation landscape section in supplement for description on an exception case). To avoid dilution of the focus of this paper, we present more detailed analysis of mutations in the Supplement information (see supplement section Mutation Landscape, mutation mapping to regions in genome and its correlations to distinct signatures such as nature of repetitive elements (Figure S3), identification of pericentromeric and heterochromatin region based on methylation signatures of H3k9me3 and H3k27me3 (Figure S4), Genome wide density plot for triple mutations (Figure S5)). Altogether, these data suggest that adaptation to SN-38 was not associated with either mutations in functional genes or changes of expression of these genes.

### Mutations result from repair of Top1-generated DSBs

To understand how hundreds of thousands of mutations in repeats and untranslated regions (see Mutation landscape section in the Supplement) could be involved in adaptation to SN-38, we proposed that they could result from DNA breaks generated by Top1. Indeed, the mutation sites strongly co-localized with sites of DNA cleavage by Top1 (Baranello *et al*, 2016) (19.8% of cases). By applying hypergeometric distribution algorithm (see Materials and Methods) we established that such co-localization is highly statistically significant (p<<e^−198^). This percent of overlap is probably a strong underestimation since experimental conditions in two studies were different, i.e. long-term generation of mutations vs short-term Top1 inhibitor treatment. Furthermore, when overlaid distribution of triple mutations along the chromosomes over distribution of Top1-cleavage sites, we observed a strong overlap of peaks (Figure 3D). Therefore, mutations take place at Top1-cleavage sites.

There are several lines of evidence that most of the mutations were generated either by HR or NHEJ repair of DSBs. 16% of the mutations in the isolates were clustered (7,510 out of 46,099 triple mutations sample), where 2-7 mutations were present within regions of up to 100 bp (we chose this length of DNA for definition of clustered mutations, see Materials and Methods, (Figure 4A & Tables S17, S18). Since random positioning predicts that mutations should be on average separated by about 10,000 bp (about 300,000 mutations per clone distributed over 3 billion base pairs of the total genome), such clustering demonstrated their non-random appearance and suggested a mechanism of their generation. In all these cases, these were loss of heterozygosity (LOH, herein referred as revert back from a mutated allele to reference allele) mutations. These LOH mutations did not result from deletions of one of the alleles (since the number of reads corresponding to these regions were similar to the average number of reads along the genome), but by copying of an allele from one chromosome to another, including copying of the entire mutation cluster (Figure 4B). Such copying could result only from the HR repair of DSBs. LOH occurred in 78% of common mutations (36,228 mutations out of 46,099), and similar fraction of LOH was found with the overall set of mutations in resistant clones (Table S19), suggesting that the majority of mutations were generated by the HR repair system.

**Figure 4.**
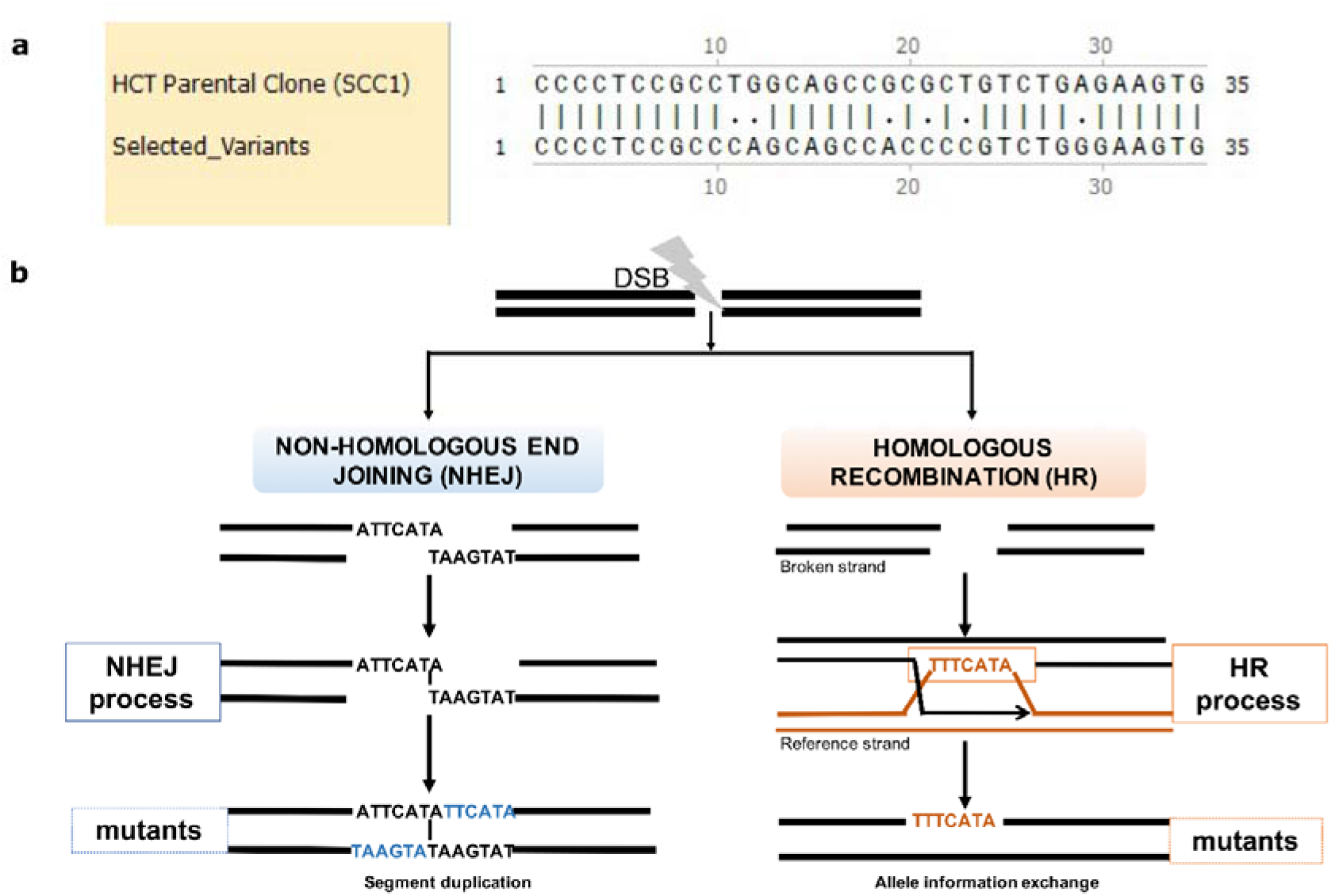
Mutations emerges in clustering patterns resulting from HR and NHEJ repair: **(a)** Example for clustering pattern of mutations compared to parental SSC1. **(b)** Sketch diagram of the mechanisms of DSB repair that leave specific mutation signatures. Real mutations found in all three resistant clones that result from NHEJ and HR repair are shown as examples. During NHEJ repair both DNA ends are either trimmed, which results in deletions, or alternatively are paired via short homology region, which results in duplication of strands on both strands (insertions) (left panel). In case of HR, an unbroken allele at the homologous chromosome can serve as template that is copied to the broken strand. Sister chromatid can also be used as template, but we do not see such events since they not generate mutations.

In the resistant clones 28% of de-novo mutations that were not LOH resulted from NHEJ (9,871 de novo common mutations, and similar fraction of NHEJ was found in the overall set of mutations). They resulted from NHEJ because in case of insertions, these mutations have a very specific signature of duplication of a neighboring region. This duplication results from pairing of broken ends at the terminal nucleotides and filling the gaps on both strands via translesion DNA synthesis (Figure 4B). This structure allows precise identification of the site of the DNA break, i.e. at the site of pairing between the duplication regions (Figure 4B) (see also Mutation landscape section and Table S20). Overall, de novo appearing InDels can be used as hallmarks of DNA breaks that were repaired via NHEJ, while LOH mutations can be used as hallmarks of DNA breaks repaired via HR.

Very importantly, a high fraction of the overlapping mutations between independent resistant clones suggests that the SN-38-inhibited Top1 generates DNA breaks at specific sites in the chromatin (possibly specific Top1 binding or activation sites), which further leads to generation of mutations upon HR or NHEJ repair of DSBs..

### Mechanism of adaptation to SN-38

The following considerations brought a framework for understanding the mechanism of adaptation. At each genome location, the parental SSC1 population could have alleles either with no mutations compared to the Reference Genome (0/0), with mutation in one allele (0/1) (heterozygosity), or both alleles (1/1) (Figure 5A, B). Sites where two alleles in SSC1 had different mutations compared to the Reference Genome (1/2) were extremely rare. Accordingly, mutations that appear in MSC clones compared to parental SSC1 could be mutations de-novo generated by NHEJ (e.g. 0/0→0/1) or loss of heterozygosity generated by HR (0/1→0/0; or 0/1→1/1) (Figure 5C, D).

**Figure 5.**
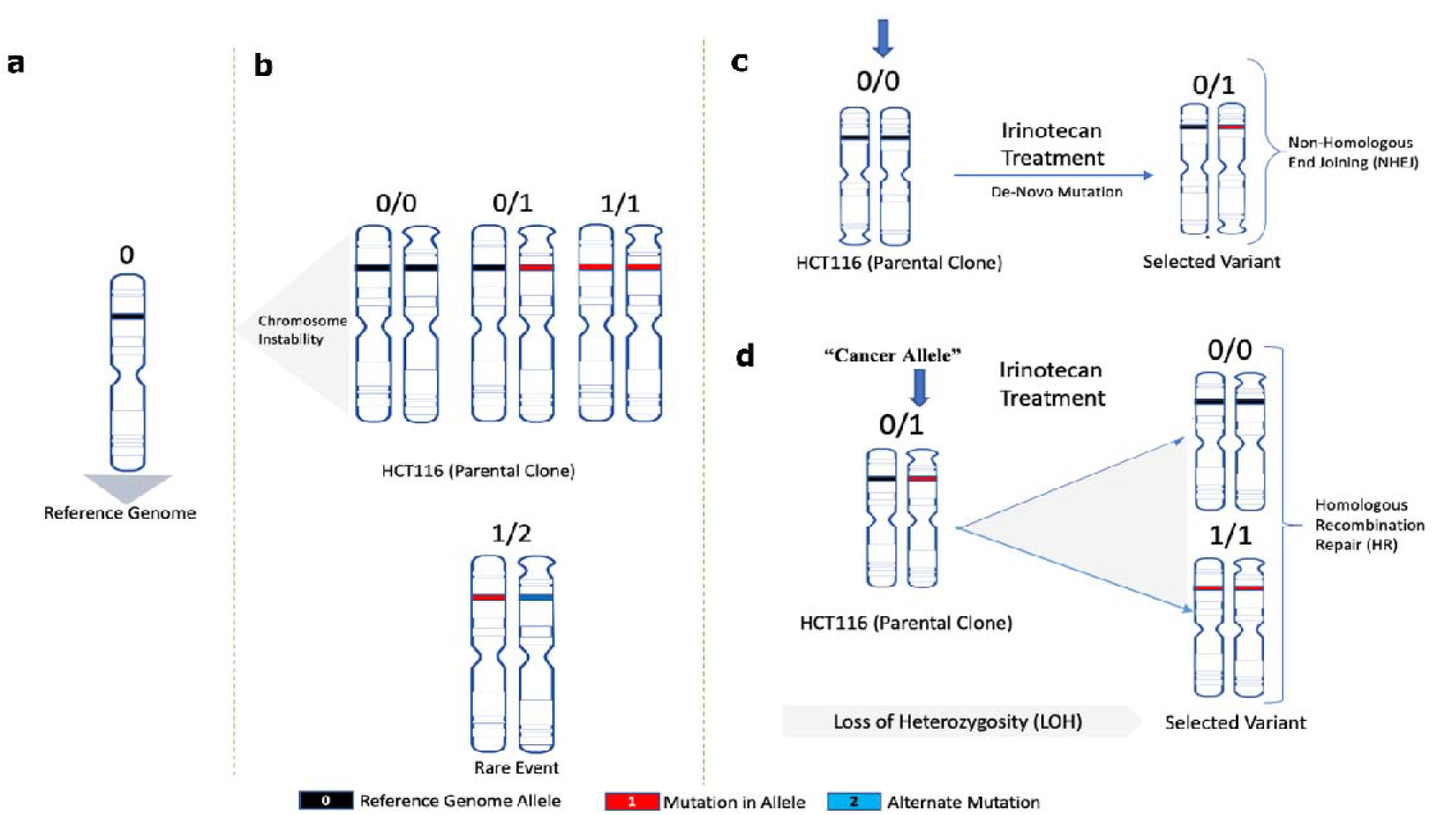
A framework model for analyses of the role of mutations in adaptation to SN-38: Schematic representation for **(a)** Normal reference genome allele, **(b)** Alleles found in parental SSC1 clone, **(c)** De-novo mutations in MSC clones after NHEJ repair, **(d)** Allele change upon loss of heterozygosity after HR repair (Loss of heterozygosity herein is referred to as the change of mutated heterozygous alleles reversed to homozygous reference in selected variants).

In sites with LOH repaired by HR, a heterozygous allele could revert to either the Reference Genome allele (0/1 to 0/0) or to the “cancer” allele seen in the parental clone SSC1 (0/1 to 1/1). A surprising key observation that led to understanding the mechanism of adaptation was that the frequency of 0/1 to 0/0 shifts was 5.44 times higher than 0/1 to 1/1 shifts (Figure 6A). 0/1 to 0/0 shift means that the allele from the Reference Genome was copied to the DSB at the “cancer” allele (an allele in SSC1 parental cells that differs from the Reference Genome), which ultimately means that the probability of double strand DNA breaks in “cancer” alleles is 5.44 time higher compared to the Reference Genome allele. The probability of breaks in “cancer” allele was even higher in the pericentromeric regions, where the ratio of breaks in Reference Genome allele to “cancer” allele was 1/20 (Fig. 5B). In the chromosome arms this ratio was 1/3.4. Therefore, surprisingly, alleles that acquired mutations in the course of cancer development were significantly more prone to Top1-induced double strand breaks than normal human genome alleles (Figure 6B).

**Figure 6.**
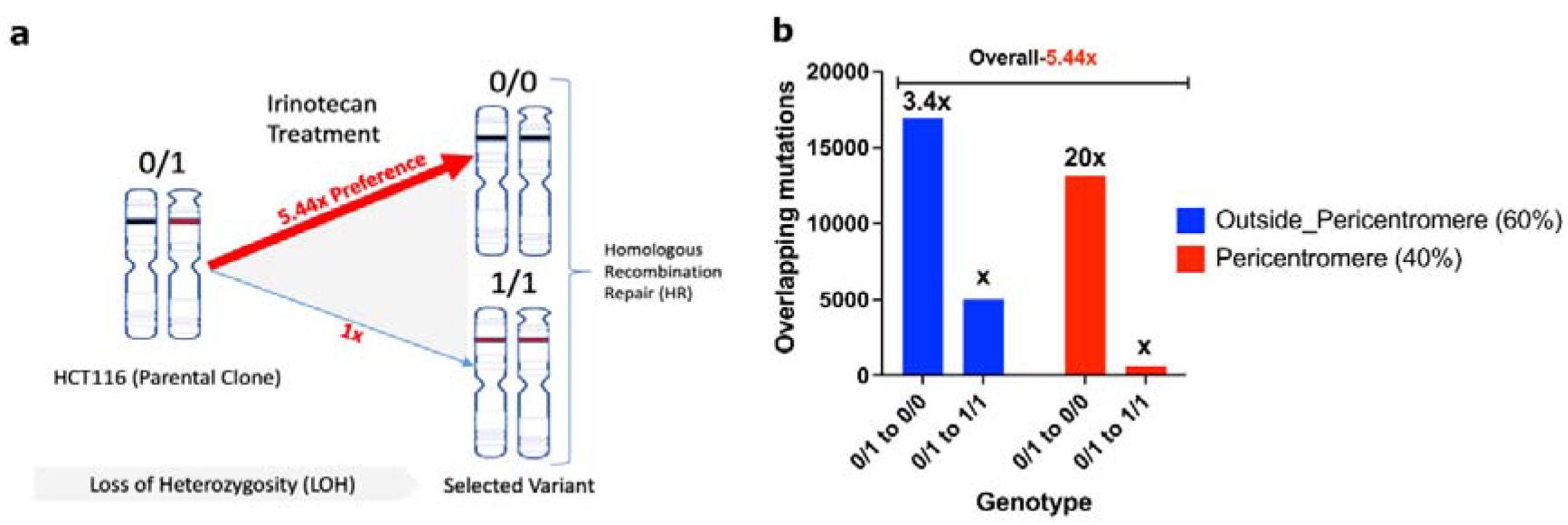
Cancer alleles are more susceptible to breaks compared to reference genome: **(a)** HR repair generated mutations revert to the reference genome allele 5.44 times more often than to the cancer allele. Therefore, cancer alleles are broken by Top1 5.44 times more often that reference genome alleles. Accordingly reverting to the reference genome alleles protects from DSBs upon consequent exposures to SN-38. **(b)** Prevalence of breaks in cancer alleles compared to reference genome alleles is lower in chromosome arms than in the pericentromeric region.

This unexpected finding provides the mechanism of gradual adaptation to SN-38. Initial exposures to SN-38 generate reversion of a number of “cancer” alleles to the Reference Genome alleles, which are more resistant to Top1-induced DSBs, which in turn creates a protective mechanism against following exposures. In other words, with each exposure there will be fewer and fewer potential Top1-cleavage sites, which will lead to stronger and stronger adaptation. This novel mechanism of adaptation to SN-38 does not involve expression of any protective proteins or mutations in them, but rather involves a high number of changes in the DNA structure that make it less prone to Top1-induced breaks.

An important consequence of this mechanism is that the cells become adapted specifically to SN-38, and do not develop resistance to other genotoxic agents. Indeed, the SN-38 resistant clones remained highly sensitive to the intercalating agent inhibitor of Top2 i.e. Doxorubicin (Figure S6).

### Adaptation associates with reduced ability of SN-38 to trigger DNA breaks

This mechanism predicts that in the process of adaptation, following multiple exposures to the same concentration of SN-38, cells should experience fewer DSBs, while the rate of repair of DSBs remains the same. To test this prediction, we took SSC7 cells that have not been drug exposed, and that underwent five cycles of exposure to 4nM of SN-38. Both populations were subjected to 4nM of SN-38 for 24 hours, and the number of γ-H2AX foci was assessed by immunofluorescence. While SN-38 exposure of naïve cells caused a dramatic increase in the number of foci, exposure of cells that were preadapted by five cycles of SN-38 treatment barely caused foci formation (Figure 7A, B). On the other hand, the rate of DSB repair (recovery of γH2AX foci) was not accelerated (Figure 7C). Importantly, lower foci formation correlated with the lack of cell death and growth arrest. Similarly, there was a lower overall number of DNA breaks in adapted cells, as judged by the comet assay (Figure 8A-E). Similar experiment was done with parental SSC1 and SN-38 adapted MSC1, MSC2 and MSC3 clone, but instead of 24 hours, exposure to the drug (40nM) was for 3, 6 and 12 hours. Figure 9A shows that overall phosphorylation of γH2AX level gradually increases in course of experiment as shown in immunoblot. As with SSC7 experiment, the drug-adapted mutants showed much fewer DSB than the parental clone at 24 hours of exposure (Figure 7A, B). Similar effects were seen upon quantification of the number of γH2AX foci in SCC1 and the mutants, representative images for 24-hour treatment are shown (Figure 9B, 9C) considering the highest difference in γH2AX levels on immunoblots (Figure 9A). Similar experiment was performed with doxorubicin to quantify the γH2AX foci after 24-hour treatment evidently, we observe foci generation in both SCC1 and SN-38 resistant mutants (Figure 9B and Figure 9D). In order to check the foci generation at lowest time point we chose 30 minutes short treatment to SCC1 and mutants and quantified the γH2AX, at this time point also we observed significant difference in the foci formed in SCC1 and mutants (Figure S7A, B). Therefore, indeed reversion of “cancer” alleles to the Reference Genome alleles appears to be associated with fewer DSBs by SN-38.

**Figure 7.**
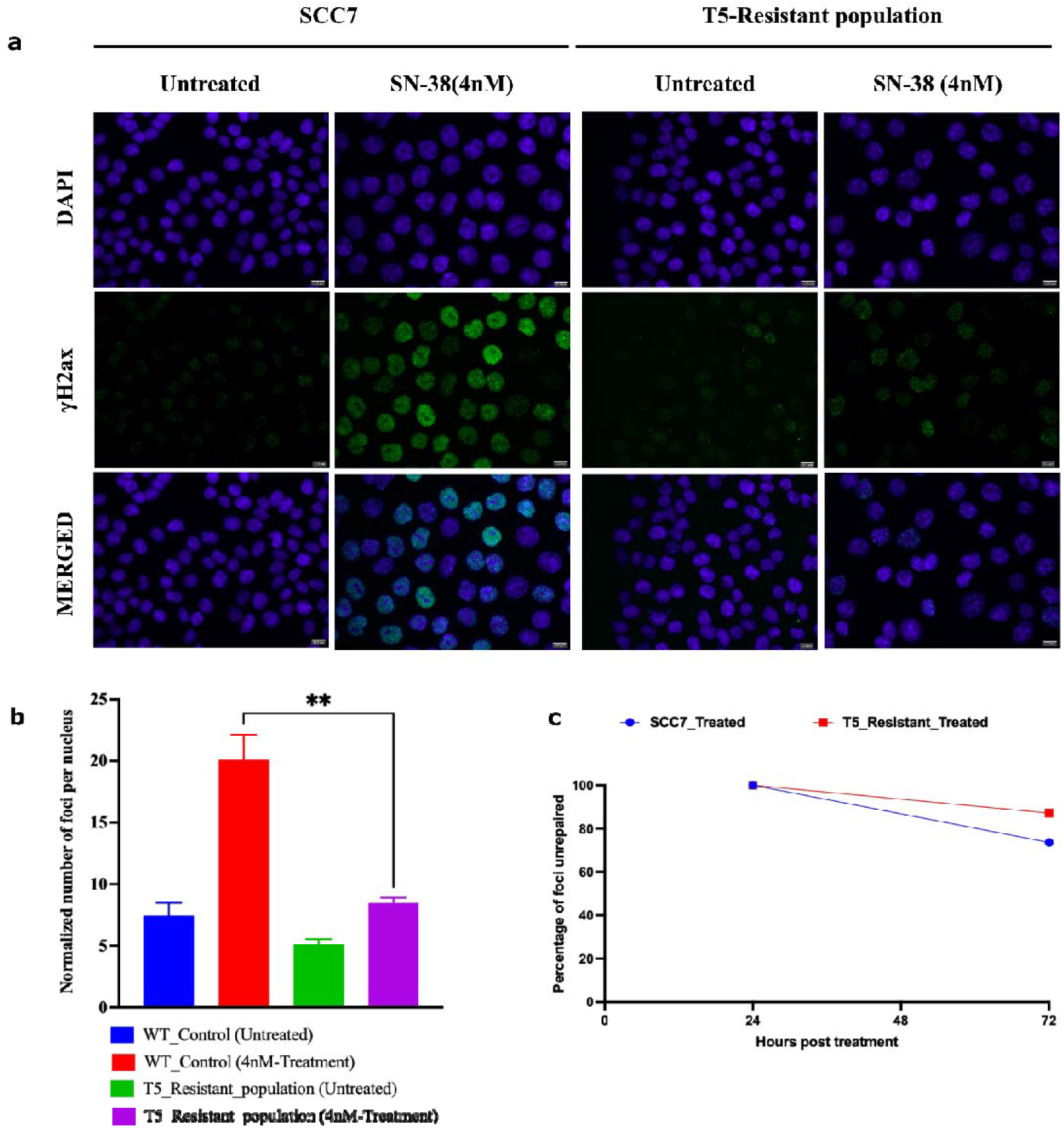
Adapted population experiences lower frequency of DSB following SN-38 exposure. **(a)** γH2AX foci in cells exposed to SN-38 (4nM for 24 hours). SCC7 cells are compared to the same clone but after five cycles of adaptation to 4nM SN-38. Experiment was conducted in biological triplicates. **(b)** Quantification of data presented in Figure 7A of the number of foci generated 24-hour post treatment, n=368 images were analyzed with the integrated software (refer material and methods). **(c)** Line plot shows the number of foci remaining after 72-hour of recovery from SN-38 (SCC7-73%, T5_resistant-87%), indicating that the rate of DSB repair is not faster in adapted cells, data represents n=431 images were analyzed using integrated software (refer material methods). Statistics were calculated using GraphPad Prism version 9.0.0, California USA. The significance of differences was determined using unpaired Welch’s correction, two-tailed t-test (**PlJ<lJ0.0021) denoted in above in figure **(b)**.

**Figure 8.**
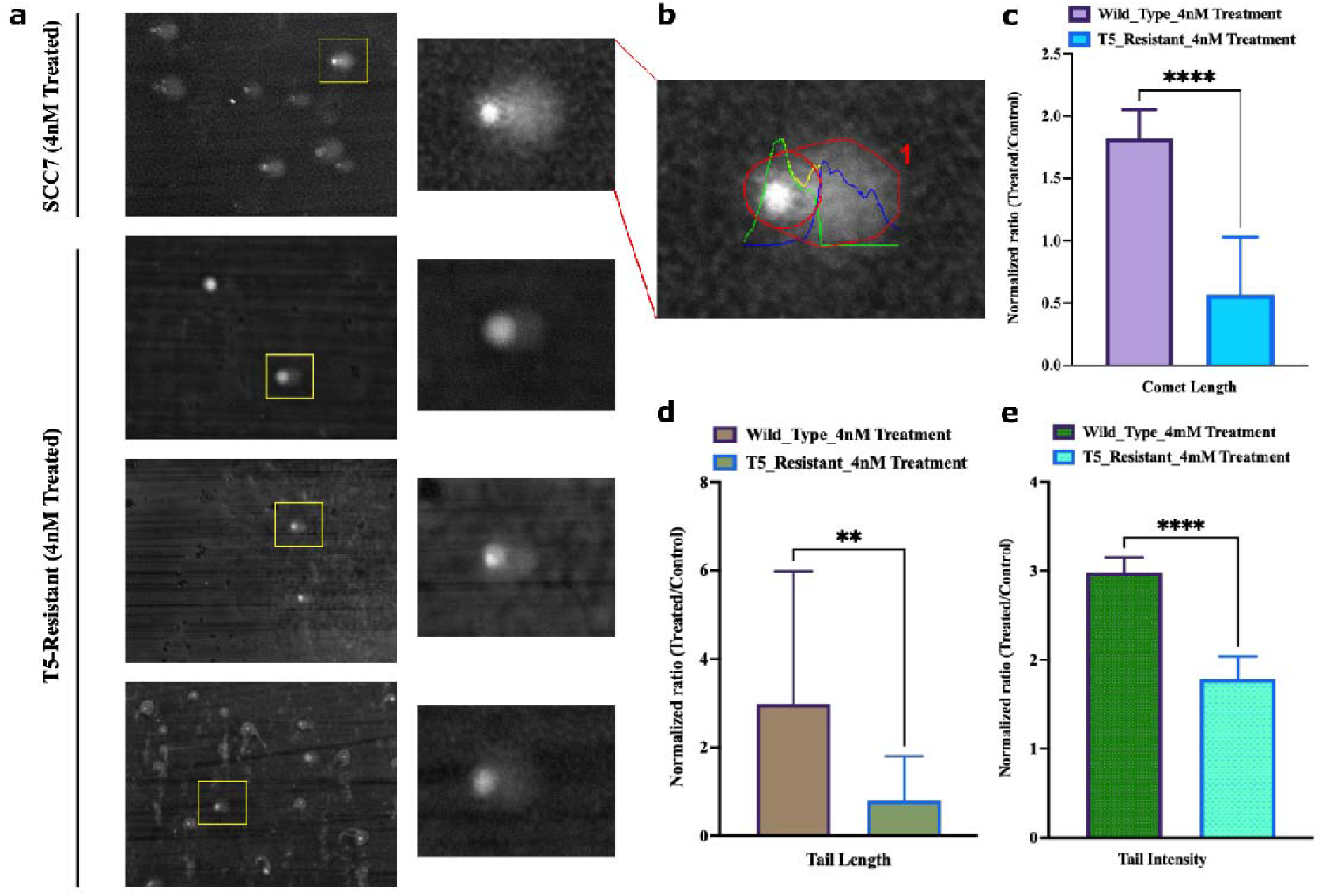
Adapted population experiences lower frequency of overall breaks following SN-38 exposure: **(a)** Representative image of the field of SCC7 cells treated with SN-38 (24 hours) and corresponding comets. Insert shows enlarged image of the comet. Lower three images with inserts show representative images of the adapted population. **(b)** Representative of quantified output image used to compute parameters for comet analysis. **(c)** Comet length and **(d)** Comet Tail length, **(e)** Intensity of DNA in tail was estimated using OpenComet v1.3.1. Bar plot represents normalized ratio of treated and control comets in SCC7 and T5-adapted cells separately. Statistics were calculated as mean of n=21 comets in each using GraphPad Prism version 9.0.0, California USA. The significance of differences was determined using unpaired Welch’s correction, two-tailed t-test (**PlJ<lJ0.0021, ****P <0.0001) denoted in above figures (C, D, E).

**Figure 9.**
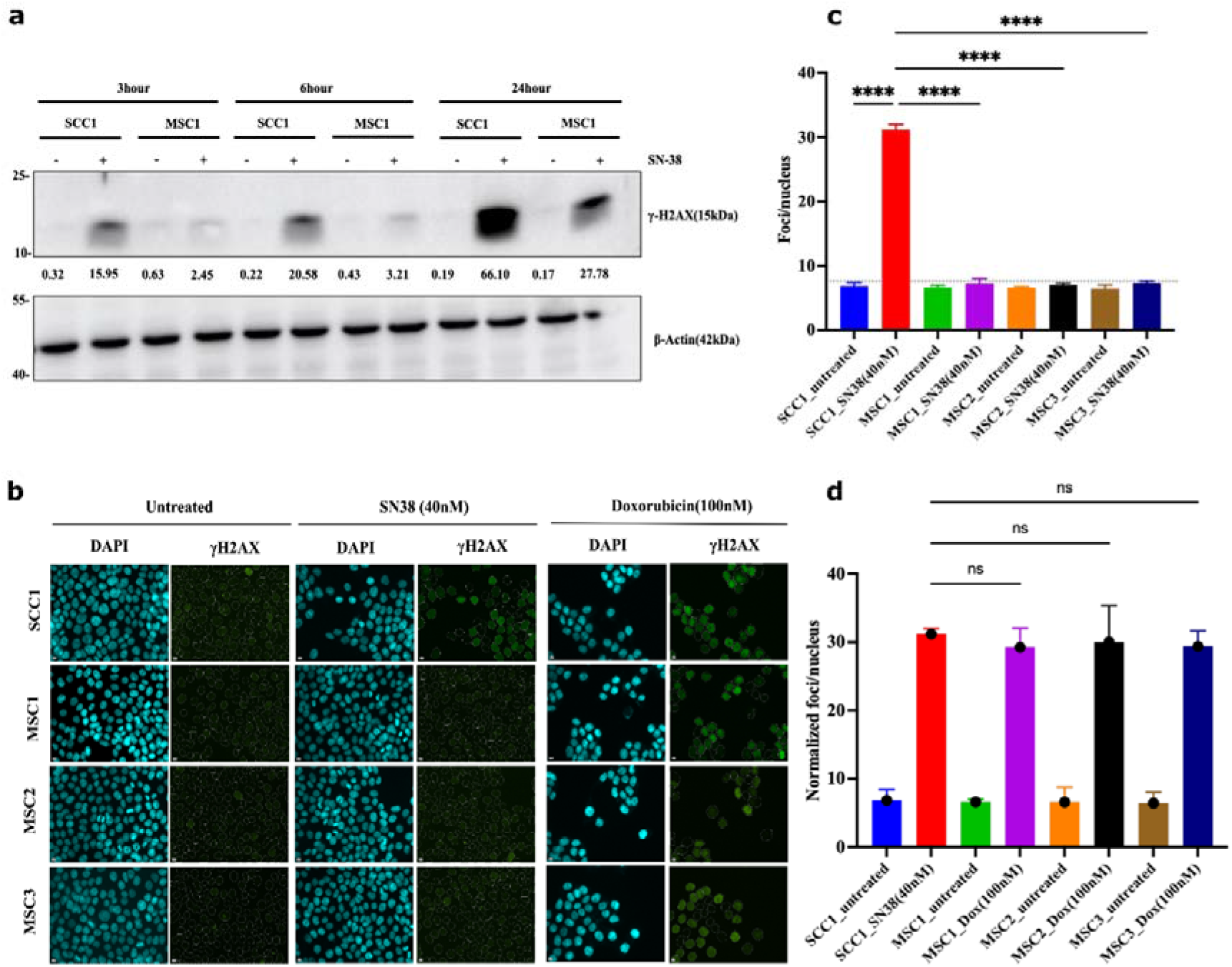
Mutants experience lower frequency of DSB following SN38 exposure: **(a)** γH2AX foci in cells exposed to SN38 (40nM for 30 min). After 3,6 and 24 hours, cells were harvested and lysed for immunoblot for quantification of phosphorylated H2AX levels as shown, experiment was conducted in biological duplicates (n=2). **(b)** SCC1 cells are compared to the mutants clones using γH2AX foci quantification for 24 hour treatment observing significant difference based on immunoblot phosphor-H2AX levels, we observe consistent increase in protein levels with highest expression for SCC1 at 24hr. Experiment was conducted in biological triplicates (n=3). **(c)** Quantification of SN-38 generated foci presented in Figure 9B showing significant decrease of foci generation in resistant mutants. n=228 images were analyzed with the integrated software (refer material and methods), for short term treatment (30 min data) and foci quantification refer Figure S7. Quantification for Doxorubicin cell sensitivity is presented in Figure S8. Statistics were calculated using GraphPad Prism version 9.0.0, California USA. The significance of differences was determined using unpaired Welch’s correction, two-tailed t-test (ns <0.1234, ****P <0.0001) denoted in above in figure (c). Pseudo-color is used in DAPI for visualization purpose. Controls served in Fig 9C and D is same as are from one experiment.

Another prediction from the suggested mechanism of resistance was that in the MSC mutants the ability of Top1 to interact with DNA is reduced compared to the parental SSC1 clone. Therefore, we sought to compare the amount of Top1 covalently bound to the chromatin following SN-38 treatment in SSC1 and MSC1 clones using DNA-Top1 adduct capturing RADAR assay. Indeed, in the presence of SN-38, in resistant mutant MSC1 the amounts of Top1-DNA adducts were significantly lower than in SSC1, further indicating that the drug adaptation associates with the loss of Top1 cleavage sites (Figure 10A,B).

**Figure 10.**
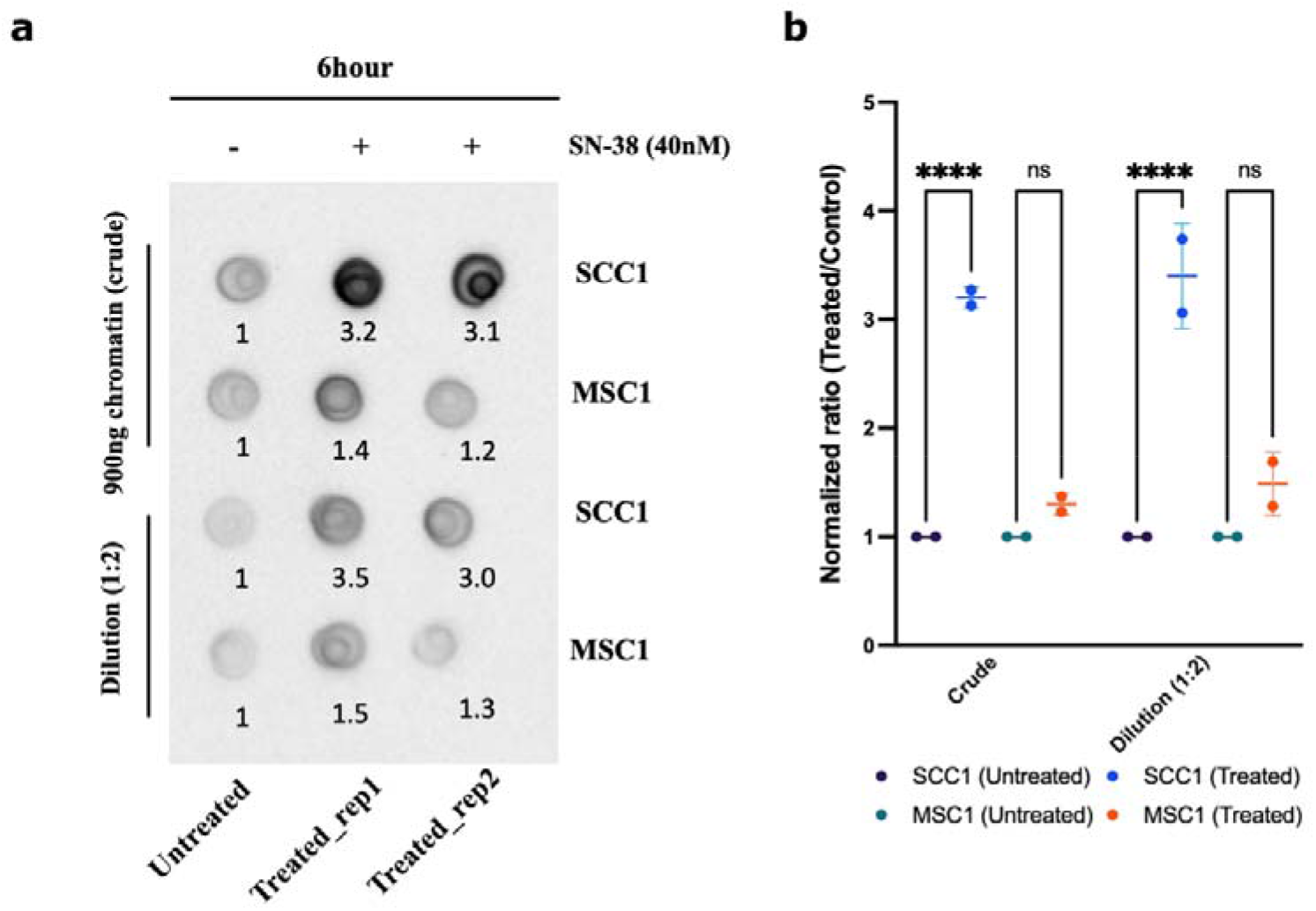
Chromatin fractionation and covalently bound Top1: Parental line SCC1 and resistant MSC1 were treated with 40nM of SN-38 for 6 hours and chromatin was isolated. Top1 covalently bound to chromatin was quantified using dot blot with Top1 antibody. **(a)** Dot blot showing the crude extract upper panel and dilution (1:2) in lower panel (For Raw blot refer raw images). **(b)** Statistics were calculated for (n=2) independent experiments using GraphPad Prism version 9.0.0, California USA. The significance of differences was determined using unpaired Welch’s correction, two-tailed t-test (ns <0.1234, ****P <0.0001) denoted.

## Discussion

Here, we addressed why the approach towards the selection of drug-resistant mutants in cancer cells requires multiple exposures to drugs and dose escalation. Such a selection scheme suggests that (a) either cells are somehow adapted to the low concentrations of drugs (e.g., via epigenetics mechanisms), and this adaptation guides further selection of the resistant mutant forms; or alternatively (b), development of drug resistance involves acquiring of a large number of mutations, each of which provides a fraction of the resistance, but gradually they accumulate and get selected in the process of drug escalation. At least with an inhibitor of Top1 irinotecan, we show that the second possibility is correct. It appeared that a very high number of mutations is generated in the process of selection of irinotecan-resistant mutants in the dose escalation experiment. Surprisingly, the absolute majority of them were in the non-coding and silenced regions of the genome, suggesting that these mutations do not affect expression or function of specific genes involved in the irinotecan resistance. Accordingly, this mechanism of adaptation is fundamentally different from previously known mechanisms related to changes in drug target, drug metabolism or transport.

During the adaptation process, we observed that survived cells acquired a senescence-like phenotype and stayed in such state without divisions for weeks and even months, until the population resumed divisions. Since senescence-like or other types of dormant states can provide drug resistance (Gordon & Nelson, 2012; Kurppa *et al*, 2020; Yuan *et al*, 2020; Duy *et al*, 2021), it is possible that acquisition of this phenotype plays a role in the ultimate survival of drug treatment.

The key to understanding the nature of the resistance was the observation that these mutations result from the repaired DSBs via the HR and NHEJ pathways. Though cuts by Top1 generate single strand breaks (SSB), these breaks can develop into DSBs (Pommier, 2013). In HR-dependent DSB repair, we observed loss of heterozygocity associated with copying of intact alleles to the allele with DSB, which allowed precise identification of the allele with DSB. Strikingly, more than 80% of DSBs took place in the alleles with mutations associated with the cancer nature of the parental cells, and accordingly, the HR repair led to restoration of the original “normal” alleles. As an example of such sequences, we found a large fraction of breaks in the polyC sequences of 30-40bp, which were present in the parental cancer cells (See Mutation landscape, Supplement Information). As a result of HR repair, these alleles were changed to alleles with interrupted polyC regions which were present in the reference human genome. Accordingly, at these sites, DSBs appear to require extended polyC, and thus an allele that has interruption of the polyC tract must have a lower probability of breaks. Therefore, reversion of extended polyC to the interrupted tract of polyC protects this site from further breaks, and thus contributes to the overall development of resistance to SN-38 (Figure S8). This observation ultimately means that upon the first exposure to the drug, a fraction of sites with high probability of Top1-induced breaks will be reverted to sites with low probability of breaks. Therefore, with each cycle of exposure to SN-38 fewer and fewer sites with high probability of DSBs will remain in the genome, which ultimately must increase the chance of survival. Indeed, we demonstrated that the number of DSB is reduced following cycles of exposure to SN-38, and this effect was associated with lower probability of breaks rather than with more efficient DNA repair. Therefore, development of resistance to SN-38 involves acquiring a large number of mutations, each of which provides a fraction of the overall resistance, which gradually accumulate and are selected in the process of drug escalation.

This adaptation mechanism that associates with gradual reduction of the number of DNA breaks by Top1, suggests lower efficiency of Top1-dependent relaxation of DNA supercoils in selected clones. Possibly, the number of mutations that provide adaptation to SN-38 may be limited by the necessity to carry out the DNA relaxation activity. Alternatively, in these clones Top2 can take over essential DNA relaxation (Figure 10, Figure S6).

Unexpectedly, the majority of the Top1-cleavage sites were in the satellite regions, especially in short repeats. This feature may explain a well-known fact that microsatellite instability in colon cancer associates with better response to the irinotecan therapy (Fallik *et al*, 2003; Benatti *et al*, 2005; Vilar *et al*, 2008). Indeed, we suggest that microsatellite instability may generate additional Top1 sites, which makes these cells more sensitive to irinotecan. Notably, the HCT116 cells that we used in the experiments have microsatellite instability (Benatti *et al*, 2005; Vilar *et al*, 2007; Chen *et al*, 2017).

Furthermore, our finding may explain a very unusual feature of irinotecan. Unlike almost any anti-cancer drug, irinotecan is not effective against stage I or II cancers and become quite effective in advanced and metastatic cancers (Köhne *et al*, 2003; Fujita *et al*, 2015; Vanhoefer *et al*, 2016; de Man *et al*, 2018; Kciuk *et al*, 2020). This puzzling feature is probably related to genetic instability in cancer, which generates additional Top1-cleavage sites during cancer evolution towards advanced stages, thus making cells more sensitive to irinotecan.

This work also illuminates novel aspects of function of Top1. Though it was reported that Top1 associates with active RNA polymerase to relieve DNA supercoils generated in the process of transcription(Baranello *et al*, 2016; Hegedüs *et al*, 2018; Singh *et al*, 2020), our data suggests that Top1 can also functions in a transcription-independent relief of supercoils since a majority of mutations was seen in heterochromatin (H3K9me3 and H3K27me3 patterns, see Figure S4). The fact that there was a very large fraction of DSBs common between the three-independent SN-38-resistant isolates indicates that there are preferable sites of breakage. This idea is reinforced by the finding that the mutation sites highly overlapped with sites of DNA cleavage by Top1 (Baranello *et al*, 2016). Possibly, these mutation sites are preferable sites of binding of Top1 or some Top1 activating factors to DNA, suggesting that Top1 or its activators have a sequence binding preference. Alternatively, Top1 may bind anywhere on the DNA and when moving together with RNApol and possibly DNApol, stalls at these regions to increase the probability of cuts. Another attractive possibility is that these repeat regions are preferable sites where SSBs are converted to DSBs.

The novel drug resistance mechanism may have interesting implications for understanding evolutionary processes. Indeed, it is possible that DSBs generated by Top1 take place predominantly at the sites of mutations that deviate from “normal” genomes. Accordingly, these DSBs can be repaired by the HR pathway, which will lead to the restoration of normal genome homozygocity (ref abstract figure). In other words, a stabilizing evolutionary selection may take place even in the absence of the selection pressure, simply as a result of the Top1 and HR repair function. Accordingly, the overall diversity of SNPs and InDels in the plurality of normal genomes has limitations that are shaped by the function of the Top1 and HR repair systems.

## Conclusions

Here we uncovered a novel mechanism of development of irinotecan resistance in colon cancer cells. We established that:

- Top1 creates cleavage sites at specific sites in DNA, mostly in the satellite regions.
- Due to the cancer evolution, cancer cells have higher number of such sites.
- Repair of Top1-generated DSB upon irinotecan treatment generates mutations at the cleavage sites that prevent interaction with Top1 upon further drug exposures.
- Accumulation of such mutations at Top1 sites over multiple drug exposures leads to reduction of Top1 binding to DNA and inability to generate toxic DSB upon irinotecan treatment.

## Acknowledgments

Additional acknowledgement to Dr. Arkadi Hesin for managing the facility for Hermes Imaging system as well as project resource management.

## Author Contributions

Conceptualization: Santosh Kumar, Michael Sherman; Methodology and pipeline design: Santosh Kumar; Investigation: Santosh Kumar, Shivani Patel, Julia Yaglom; Bioinformatics analysis: Santosh Kumar, Valid Gahramanov, Shivani Patel; Suggestions in data analysis: Ivan Alexandrov, Gabi Gerlitz, Mali Salmon-Divon, Lukasz Kaczmarczyk; Supervision: Michael Sherman; Writing-original draft: Michael Sherman, Santosh Kumar; Organizing manuscript and visualization of data: Santosh Kumar; Review and approval: All the contributors.

## Ethics approval and consent to participate

Not applicable

## Consent for publication

Not applicable

## Data Availability Statement

Data such as raw FASTQ files for human whole genome sequencing has been submitted to publicly accessible database SRA under accession number PRJNA738674. Processed VCF showing identical mutations among all resistant subclones compared to parental line (available in supplement information Table S21) is included in this paper. Full list of genes from transcriptome analysis and shRNA screens are available in this paper. Raw data and files after post analysis for transcriptome analysis is deposited and is available at GEO data base with accession number GSE189366. Analyzed data is presented in supplement table at relevant section. **Already published elsewhere used in analysis:** These data were referred from literature; GSE57628, mapping of Top-1 binding and cleavage sites reported for HCT116 cells, Summary of epigenome ENCSR309SGV (ENCODE database that present the summary of various methylation signatures such as H3K9me3 and H3K27me3 for genome of HCT116). All the codes essentially used in analysis of sequencing data, either human whole genome, transcriptome analysis or shRNA screens has been submitted to public domain and can be accessed through Github (https://github.com/santoshbiowarrior333/Irinotecan_resistance).

## Supplementary Materials

Supplementary Data are available at NAR online, additionally “Data_S1.xlsx” contains all the supplementary tables and metadata.

## Supplementary section

### Mutational landscape in SN-38 resistant clones

As noted in the main text, mutations in coding and regulatory regions did not affect genes involved in cell sensitivity to SN-38. The only exception was a mutation in the MMS22L gene that normally works in the base excision repair, but also plays a role in the DSB repair (Piwko *et al*, 2011; Saredi *et al*, 2016). However, this mutation was a 1 nucleotide insertion, which most likely inactivated the MMS22L gene. In such a scenario, we expect to see sensitization rather than protection from DSBs. Accordingly, it is unlikely that this mutation associates with adaptation to irinotecan (SN-38).

To understand the relevance of the mutations to the adaptation process, we manually identified the precise positions of breaks in 1,000 out of 2,829 NHEJ-repaired common mutation sites and analyzed if there are any shared features in sequences of the sites. About 80% of these breaks took place either within or at the edge of 1-4 bp repeat clusters, and the length of these repeats usually was within the range of 5-30bp (see examples of such repeats in Table S17, Table S18). Though we could not define the breaking points in HR repaired sites precisely, we observed that 75% of these sites are in similar repeats as NHEJ breaking points, and therefore assumed that breaks in these sites occurred in or at the edges of such repeats, as with NHEJ. Overall UCSC analysis showed that the mutations were mostly located in the repetitive elements such as microsatellites (Table S22), simple repeats (Table S23) and satellites (Table S24 for mutations in chromosomal arms and supplement (Table S25) for mutations in pericentromeric genomic locations). Parallel RepeatMasker analysis supports these findings (Figure S3A, Table S26 and S27 for detailed data on repetitive elements and mutation correlation) where specific satellites such as alpha satellites and human satellite-II is shown to have majority of common mutations (Fig S3B). In other words, repetitive elements accommodate majority of all mutations, including common mutations between the clones, during the adaptation process. Upon analysis of distribution of mutations along the chromosomes, we found that there was a disproportionally around 40% of the total mutations densely populated regions adjusted to centromere (up to 5Mb depending on the chromosome) at both sides (Figure S4), which we referred to as “pericentromeric” areas. This region is identified generally by signatures of H3K9me3 and H3K27me3 followed by richness of repetitive elements such as satellites and simple repeats (Figure S4A, B and Fig S5). As a representative example, Figure S4A shows correlation between pericentromeric locations of common mutations on chromosome-1, pericentromeric chromatin modification signatures, and locations of satellites. For comparison, Fig. S4B shows these distributions and parameters in a chromosomal arm of same chromosome-1. Whereas, Fig. S5 visualizes the overall distribution of common mutations in genome and show its specific enriched presence in centromere and pericentromeric regions.

A very common sequence at the breaking points in the chromosome arms was a polyC stretch (25% of all mutation sites in the arms). The length of these polyC sequences associated with breaking points usually were within the range of 20-40bp. To our surprise, in 100% cases of LOH in these regions, original long polyC was substituted with polyC stretch that was interrupted by multiple SNPs (Figure S8). The fact that these breaks were generated in response to Top1 inhibition by SN-38 suggests that Top1 either binds to such sites specifically, or stalls there in the process of its function, or is activated at these sites.

### Supplement figures

**Figure S1.**
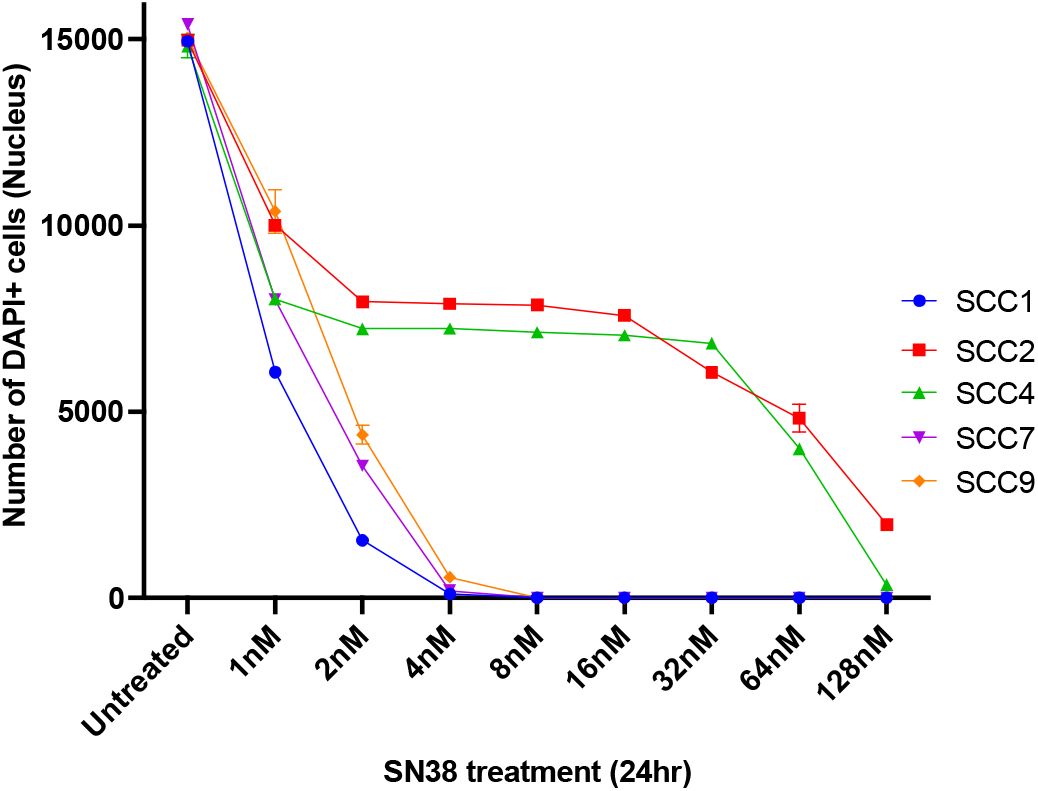
Sensitivity of the single cell clones to SN-38: Isolated clones were grown individually on 24 well plates followed by treatment with several concentrations of SN-38 and were incubated for different time durations for assessment of cell death. After incubation, dead cells were washed and attached cells were fixed, followed by DAPI staining. Using Hermes imaging system, images (n=244) were taken for each well. Experiment was conducted in biological triplicates (n=3) to quantify the cell cytotoxicity.

**Figure S2.**
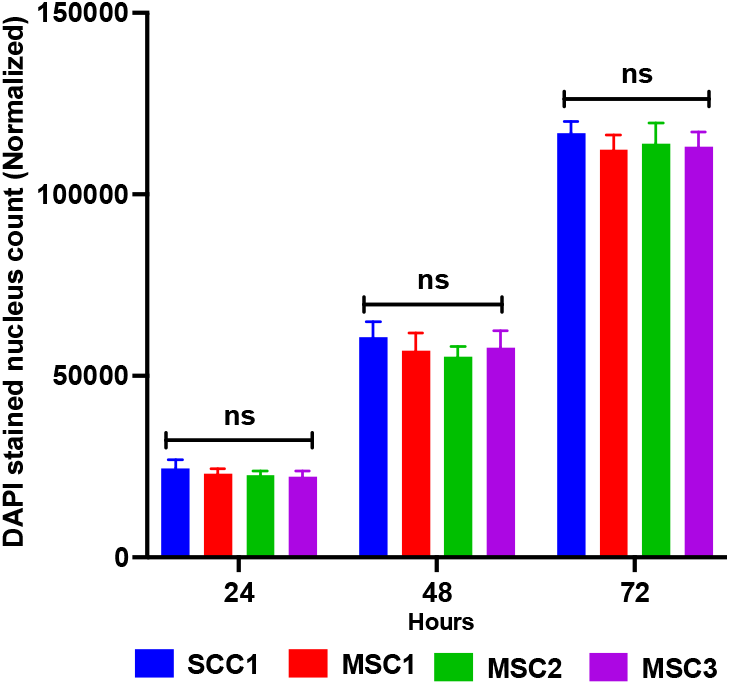
Growth rate of parental clone and resistant mutants: Equivalent number of parental clone population (SCC1) and resistant mutants were plated and kept for normal growing. At time 24-hour interval, cells were fixed and stained by DAPI. Imaging was performed by Hermes imaging system where n=268 images were taken per well; experiment was performed with six biological replicates (n=6). Data shows non-significant difference in increase of cell population among parental clone or mutants indicating the growth rate of parental and mutant are similar in culture. Statistics were calculated using GraphPad Prism version 9.0.0, California USA. The significance of differences was determined by Ordinary one-way ANOVA, error bar represents standard deviation (computed *P=*0.2009 is “ns”) as denoted indicating that there was no significant difference at same time points in SCC1 or resistant mutant growth.

**Figure S3.**
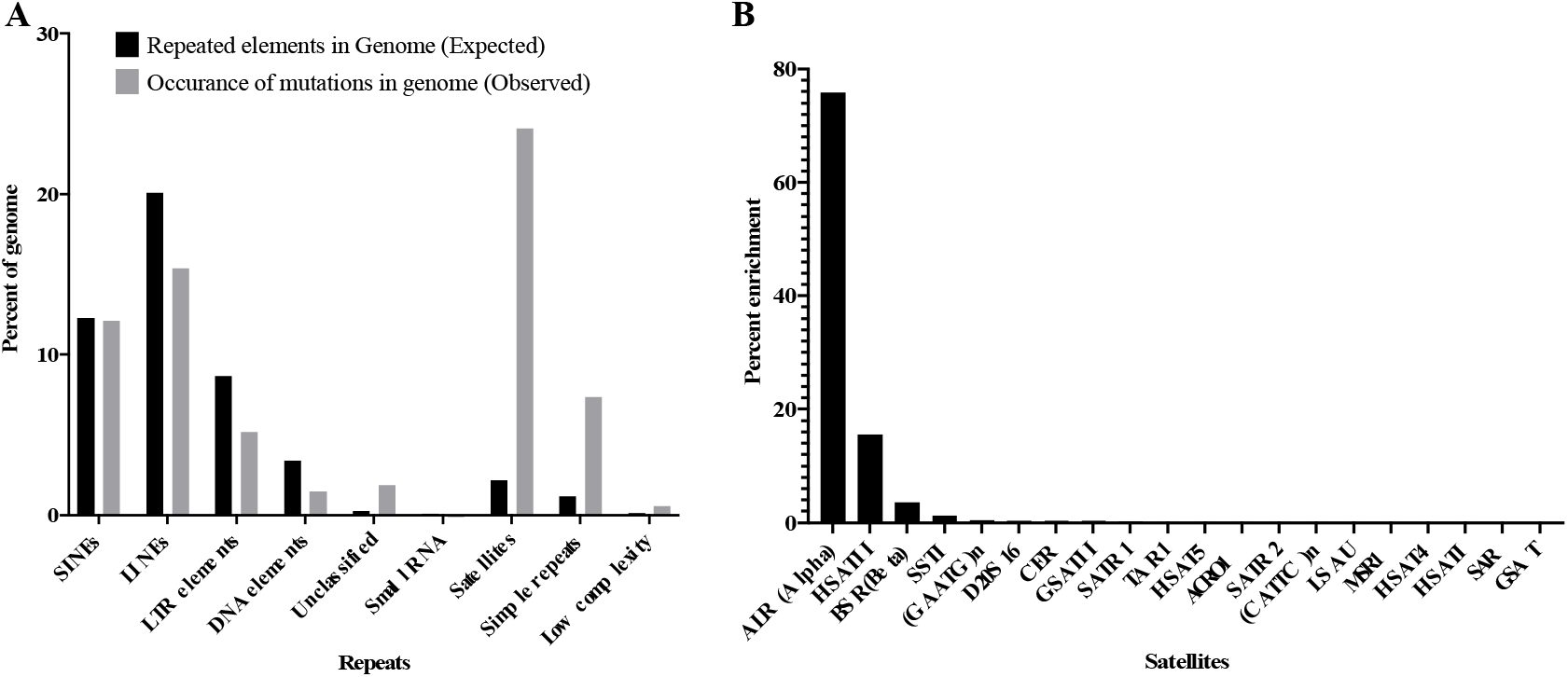
**(A)** Distribution of repeated elements in human genome (Expected) and fraction of mutations in MSC clones found in these repeats. common mutation sites were taken randomly (100bp upstream and downstream of SNP’s) and run through the RepeatMasker (v4.1.0) using nhmmscan version 3.3.1 (Jul 2020). The presence of mutations in SINE corresponds to the presence of SINE in the genome, indicating lack of the enrichment. Strong enrichment was found in satellites and simple repeats, **(B)** In-depth analysis for classification of satellites shows vast majority of common mutations are present in ALR and human satellite II followed by BSR type in all three clones.

**Figure S4.**
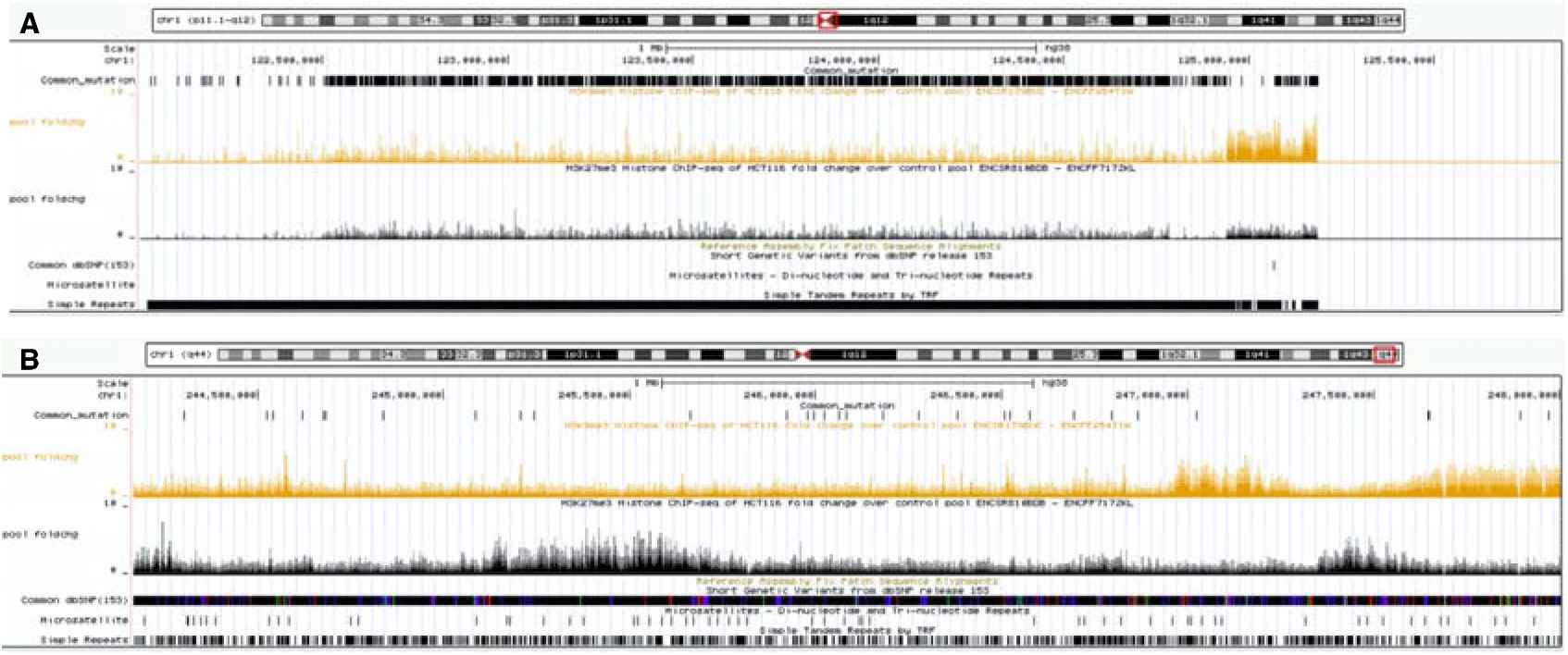
Representative example for pericentromeric and chromosomal arm: **(A)** plot shows correlation between common mutations in “pericentromeric region” marked by H3K9me3 and H3K27me3, where mutations are densely accommodated and covered by simple repeats. Mutations in these genomic locations are not common dbSNP’s and is accommodated by fewer microsatellites, **(B)** shows arm portion of chromosome with comparatively much fewer common mutations and scattered simple repeats in sliding window UCSC browser.

**Figure S5.**
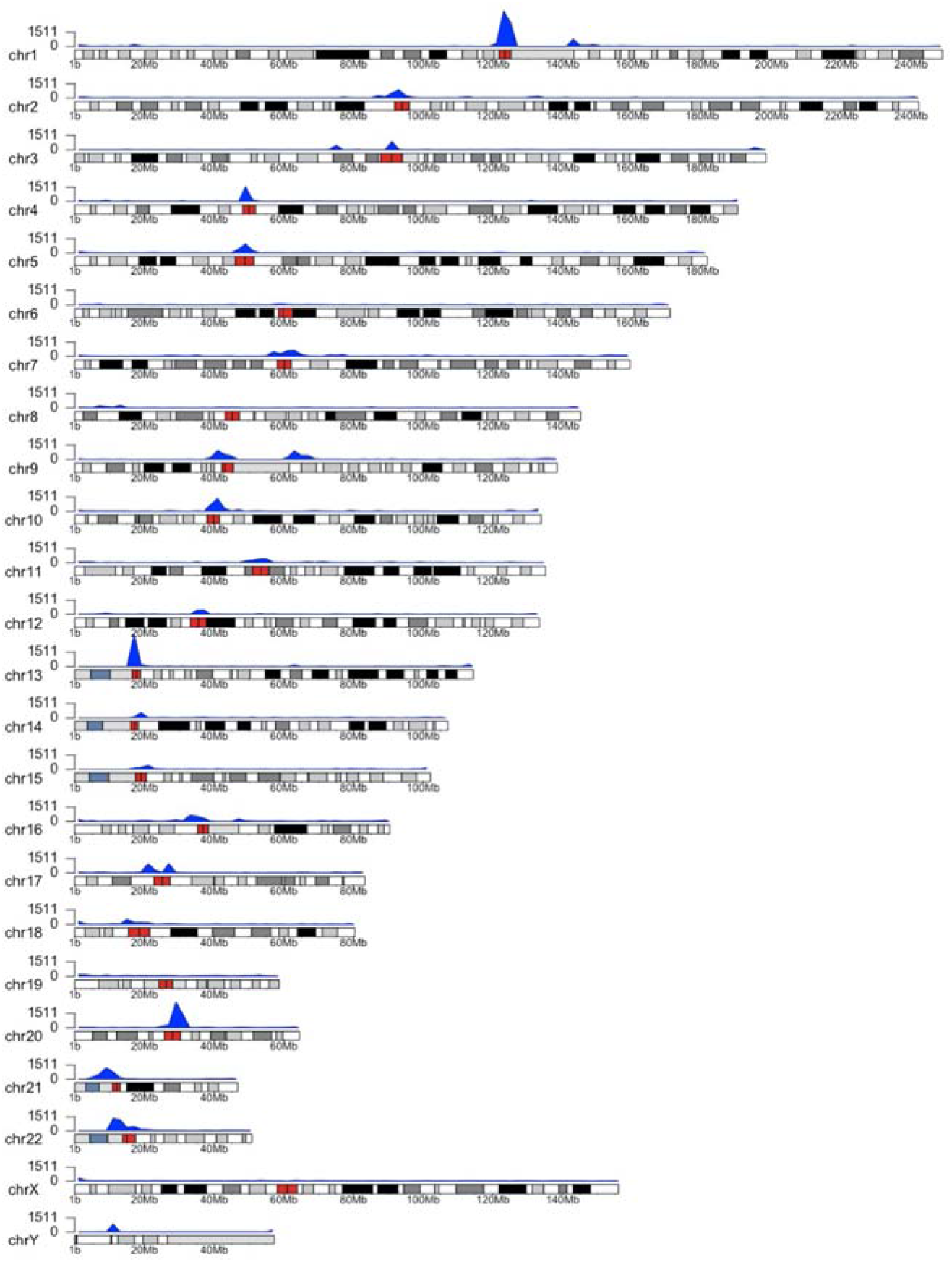
Disproportionate distribution of common mutations in genome: Karyoplot showing enrichment of mutations (blue peaks) in pericentromeric region (marked with red on cytoband) where genomic positions have been plotted against SNP density in corresponding regions using R package (karyoploteR 1.18.0).

**Figure S6.**
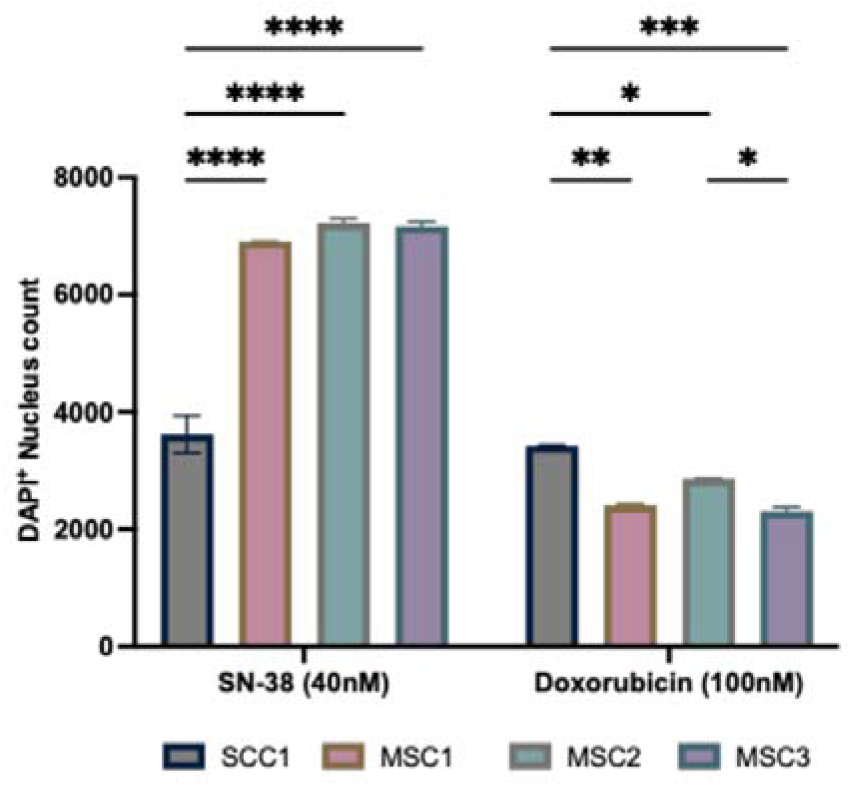
SN-38 resistant mutants retain sensitivity to doxorubicin: Cells were treated with SN-38(40nM) or doxorubicin (100nM) for 24 hours or left untreated. After 24 hours, cells were washed to remove dead cells and fixed, stained by DAPI. Experiment was conducted in biological triplicates (n=3). Quantification of data presented is the number of DAPI^+^ nucleus count after treatment, n=183 images were analyzed with the integrated software (refer material and methods). Statistics were calculated using GraphPad Prism version 9.0.0, California USA. The significance of differences was determined using Two-way NAOVA (*PL<L0.0332, **PL<L0.0021, ***PL<L0.0002, ****P <0.0001) denoted.

**Figure S7.**
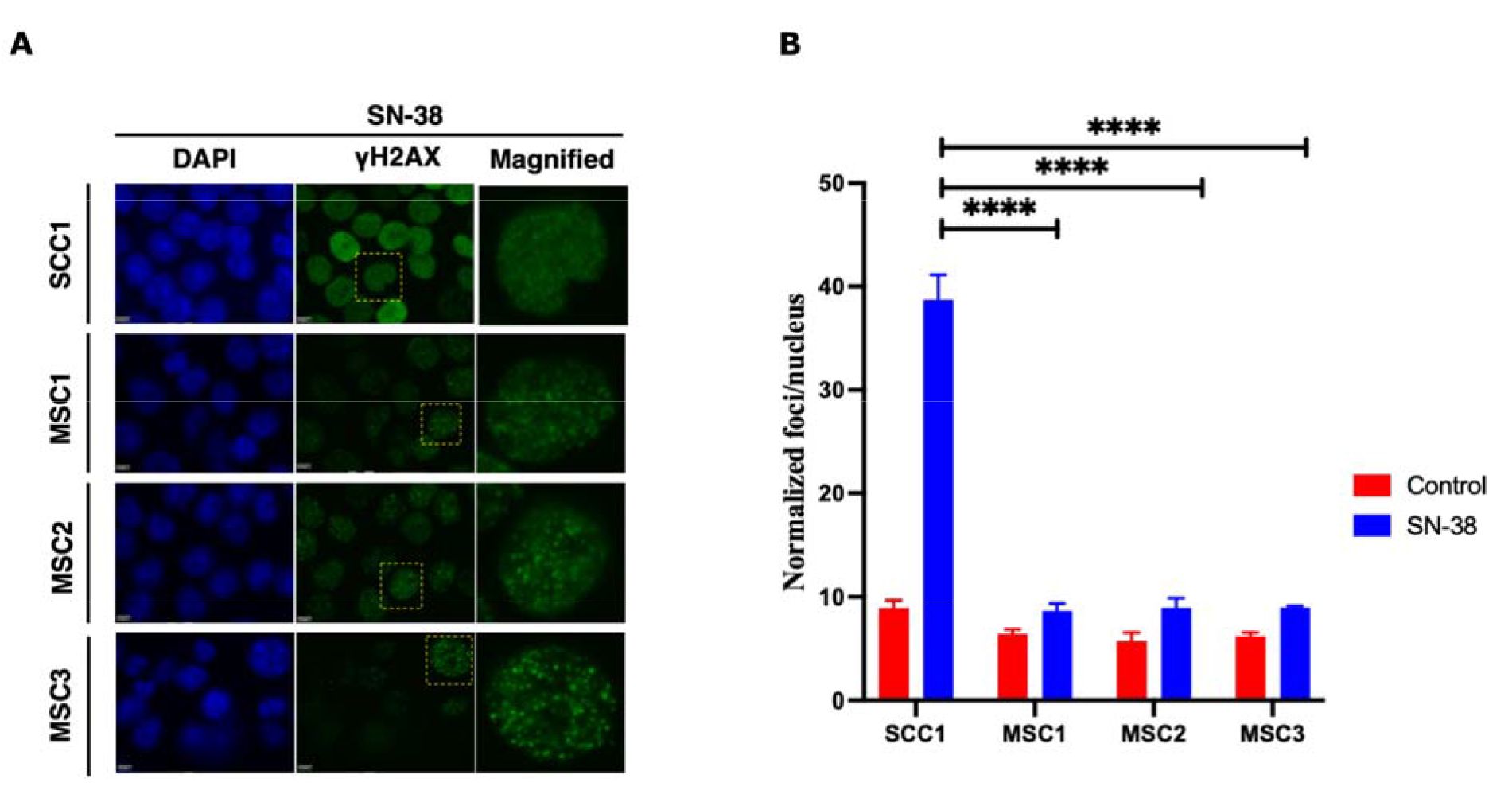
Resistant mutant experiences lower frequency of DSB following SN-38 exposure to short treatment. **(A)** γH2AX foci in cells exposed to SN-38 (40nM for 30min). SCC1 cells are compared to the mutants after very short treatment. SCC1 experiences significantly higher DSB compared to resistant clones, for long exposure data refer (Figure 9A-C). Experiment was conducted in biological triplicates. **(B)** Quantification of data presented in Fig. S9A of the number of foci generated 30-min post treatment, n=168 images captured at 40x, were analyzed with the integrated software (refer material and methods). Statistics were calculated using GraphPad Prism version 9.0.0, California USA. The significance of differences was determined using unpaired Welch’s correction, two-tailed t-test (****P <0.0001) denoted in above in figure (B).

**Figure S8.**
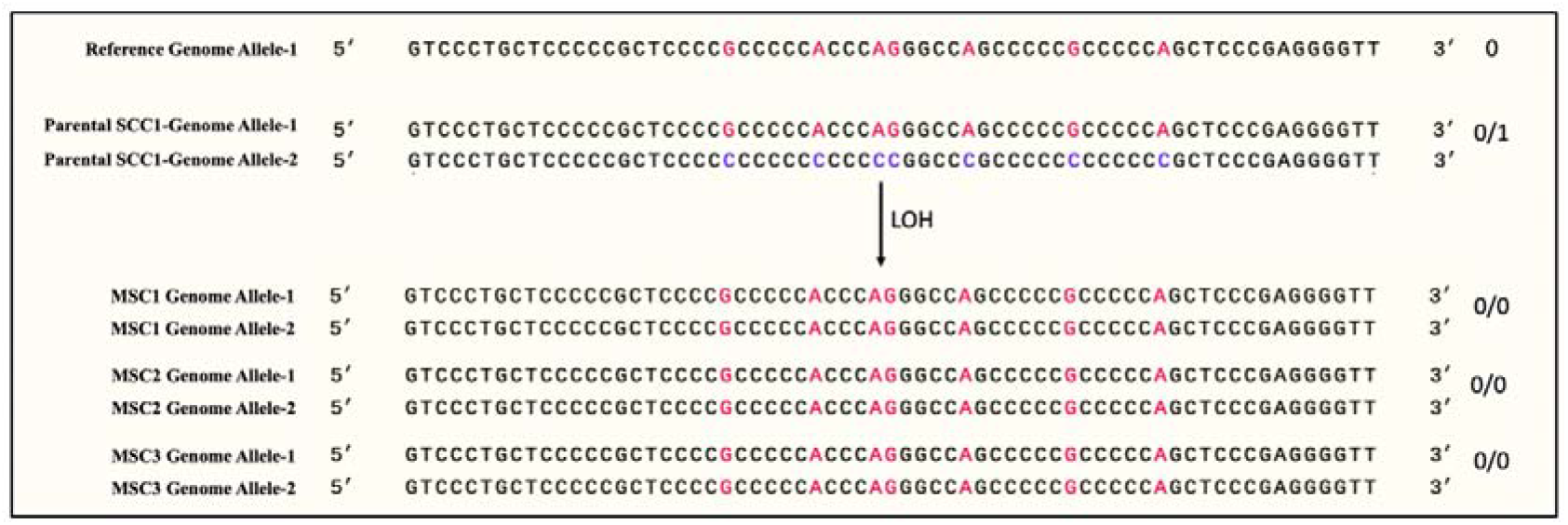
Representative example mutations in polyC sequences. Comparative alignment of a polyC sequence in SSC1 with reference genome (hg38) and resistant sub-clones. Allele-1 and Allele-2 represent two alleles in homologous chromosomes. Sequence marked by (both red) alleles represents homozygosity (0/0) as shown in reference genome strand and resistant sub-clones (top). Alleles (red and blue) in parental line represent heterozygosity (0/1) with “Cs” base. After the SN-38 adaptation, generated mutations lead to LOH, so that the polyC tract became interrupted as in normal reference genome alleles.

